# Predicting genotype-specific gene regulatory networks

**DOI:** 10.1101/2021.01.18.427134

**Authors:** Deborah Weighill, Marouen Ben Guebila, Kimberly Glass, John Quackenbush, John Platig

**Affiliations:** Harvard T.H. Chan School of Public Health, Boston, MA 02115, USA; Channing Division of Network Medicine, Brigham and Women’s Hospital, Boston, MA 02115, USA; Harvard Medical School, Boston, MA 02115, USA

**Author notes:** Current address: Lineberger Comprehensive Cancer Center, University of North Carolina at Chapel Hill, Chapel Hill, NC, USA.

## Abstract

Understanding how each person’s unique genotype influences their individual patterns of gene regulation has the potential to improve our understanding of human health and development and to refine genotype-specific disease risk assessments and treatments. However, the effects of genetic variants are not typically considered when constructing gene regulatory networks, despite the fact that many disease-associated genetic variants are thought to have regulatory effects, including the disruption of transcription factor (TF) binding. We developed EGRET (Estimating the Genetic Regulatory Effect on TFs), which infers a genotype-specific gene regulatory network (GRN) for each individual in a study population. EGRET begins by constructing a genotype-informed TF-gene prior network derived using TF motif predictions, eQTL data, individual genotypes, and the predicted effects of genetic variants on TF binding. It then uses message passing to integrate this prior network with gene expression and TF protein-protein interaction data to produce a refined, genotype-specific regulatory network. We used EGRET to infer GRNs for two blood-derived cell lines and identified genotype-associated, cell-line specific regulatory differences that we subsequently validated using allele-specific expression, chromatin accessibility QTLs, and differential ChIP-seq TF binding. We also inferred EGRET GRNs for three cell types from each of 119 individuals and identified cell type-specific regulatory differences associated with diseases related to those cell types. EGRET is, to our knowledge, the first method that infers networks that reflect individual genetic variation in a way that provides insight into genetic regulatory associations that drive complex phenotypes.

EGRET is available through the Network Zoo R package (netZooR v0.9; netzoo.github.io).

## 1 Introduction

The mechanisms by which disease-associated genetic variants exert their effects on phenotype remains an unanswered question. Genome-wide association studies (GWAS) have found that phenotype-associated genetic variants typically have modest effect sizes, generally lie outside of coding regions, and likely influence gene regulation [1, 2, 3, 4]. These findings are consistent with the observation that genetic variation in transcription factor (TF) binding sites and their flanking regions explains a substantial amount of the heritability of many diseases and complex traits [5]. What is lacking are methods that can effectively predict genotype-specific TF regulatory network structure, helping to explain the genotype-phenotype link at the level of the individual.

EGRET (Estimating the Genetic Regulatory Effect on TFs) is a method built on a simple premise: genetic variants that affect both transcription factor binding in a gene’s regulatory region and the expression of that gene should produce an alteration in an individual’s gene regulatory network (GRN). More specifically, such a variant should alter the edge weight connecting a TF and its target gene in a specific individual’s regulatory network if that person carries the variant. EGRET begins with an initial guess of TF-to-gene edges using TF binding motifs identified within the human reference genome. For variants lying within TF binding motifs in a gene’s promoter, EGRET determines whether the variants are likely to influence the binding of the TF and the expression of the gene. If a variant meets these criteria, then the initial estimate of the TF-to-gene edge is modified. EGRET then uses message passing [6] to integrate these modified edges with TF-TF interaction data and population-level gene expression information to construct individual-specific GRNs where the weight of an edge connecting a TF to a gene reflects the confidence that the TF regulates that gene.

We validated EGRET in two ways. First, we inferred and compared networks for two genotyped cell lines and found that predicted genotype-specific, differentially regulated genomic regions were enriched for genotype-affected chromatin accessibility, allele-specific expression, and differential TF binding as determined by ChIP-seq; each provided independent evidence that EGRET networks capture biologically relevant genetic disruptions in gene regulation. Second, we used EGRET to infer 357 individual and cell-type specific GRNs (three cell types, 119 individuals) and showed that EGRET networks captured cell-type specific, genetically influenced regulatory disruptions in relevant disease processes. Notably, these disease-related regulatory disruptions affected network modularity in different cell types, indicating that the effects of genetic variants extended beyond differential regulation of genes to the alteration of higher-order regulatory processes through changes in network structure.

These results show that EGRET can identify genetic changes that alter cellular functions and the causal role played by disease-associated variants. Because the only individual-specific data EGRET requires is genotype data, it can be used to understand genetic effects in many cohorts with SNP chip or whole genome sequencing data, such as TOPMED [7] and the UK biobank [8].

## 2 EGRET - Estimating the Genetic Regulatory Effects on TFs

### 2.1 The EGRET algorithm

EGRET uses several sources of information to capture the impact of genetic variants on TF-to-gene regulatory relationships and construct individual-specific GRNs (Tables 1 and S1, Figures 1A and S1). The first is a TF-to-gene reference motif prior network *M* derived, for example, from motif scans of a reference genome ([9], Supplementary Note S1) that estimates which TFs bind to promoter regions to regulate target gene expression. Second, EGRET requires eQTL data either from the study population or from a public database from the cell type of interest. Third, EGRET uses genotype information in the form of genetic variants of the individual(s) for which GRNs are being constructed, and fourth, EGRET uses predictions of the effect of these genetic variants on TF binding (Supplementary Note S2) to modify the reference motif prior *M*, producing a genotype-specific “EGRET prior” *E*.

**Table 1:** Data types and sources used as input to EGRET. Note that the application of EGRET to the 119 Yoruba individuals uses the same motif and PPI priors as used in the cell line analysis.

| Data type | Tissue/Genotype | Source |
| --- | --- | --- |
| Genotype cell line comparison |  |  |
| eQTLs | LCL | https://gtexportal.org/home/datasets [11] |
| Gene expression | LCL | https://gtexportal.org/home/datasets [11] |
| Genotype | GM12878 | https://www.illumina.com/platinumgenomes.html [12] |
| Genotype | K562 | https://www.encodeproject.org/files/ENCFF538YDL/ [13] |
| ChIP-seq | GM12878, K562 | http://pedagogix-tagc.univ-mrs.fr/remap/ [14] |
| PPI | N/A | https://sites.google.com/a/channing.harvard.edu/kimberlyglass/home [15] |
| Motif | N/A | constructed using FIMO [9] |
| Population application: 119 Yoruba individuals |  |  |
| eQTLs | LCL | http://eqtl.uchicago.edu/jointLCL/ [17] |
| eQTLs | iPSC/iPSC-CM | http://eqtl.uchicago.edu/yri_ipsc/ [17] |
| Gene expression | LCL | http://eqtl.uchicago.edu/jointLCL/ [17] |
| Gene expression | iPSC/iPSC-CM | https://www.ncbi.nlm.nih.gov/geo/query/acc.cgi?acc=GSE107654 [17] |
| Genotype | Yoruba individuals | http://eqtl.uchicago.edu/yri_ipsc/ [17] |

**Figure 1:**
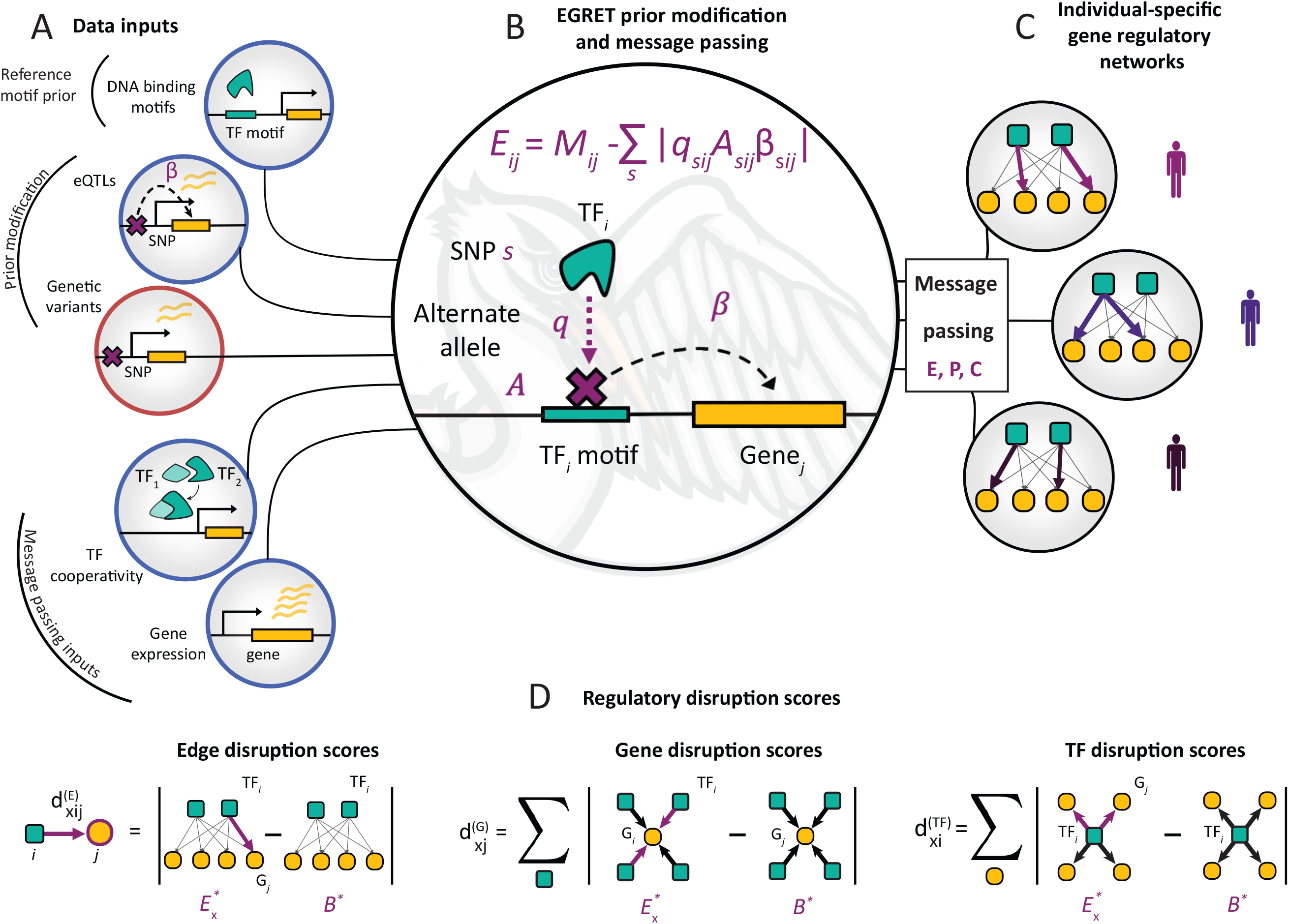
EGRET integrates multiple data types to construct individual-specific GRNs. **(A)** EGRET takes as input several data types (population-level inputs circled in blue, individual-specific inputs circled in orange) to construct individual-specific GRNs: an initial estimate of the binding locations of TFs in the form of a reference motif prior (*M_ij_*), the beta values of eQTL associations between “eSNPs” and “eGenes” (*β*), the genetic variants (*s*) harbored by the individual in question, PPI data as an estimate of TF-TF co-operativity (*P*), and gene expression to estimate a gene co-expression matrix (*C*). **(B)** An individual’s genetic variants are used to modify the reference motif prior to produce an individual-specific EGRET prior (*E*) by penalizing motif-gene connections in which that individual carries a variant allele (*A*) in the relevant promoter-region motif such that the variant is an eQTL for the adjacent gene (*β*) and the variant is predicted by QBiC to affect TF binding at that location (*q*). **(C)** Message passing is used to integrate the co-expression (*C*) and PPI (*P*) networks with the EGRET prior (*E*) resulting in a final, unique GRN per individual (*E*\*). **(D)** Regulatory disruption scores can be calculated to quantify the extent to which an edge or node in the network is disrupted by variants. Edge disruption scores are calculated by subtracting a genotype-agnostic baseline network (*B*\*) from the individual’s EGRET network and taking the absolute value. TF or gene disruption scores are calculated taking the sum of the edge disruption scores around the TF or gene in question.

Specifically, for each input genotype, EGRET selects SNPs (*A* in Figure 1B) that (1) are within motif-based TF binding sites in the promoter regions of genes, and (2) have a statistically significant eQTL association (*β* in Figure 1B) with the expression of the adjacent gene. EGRET then uses QBiC [10] to identify SNPs within TF motifs that significantly affect TF binding (*q* in Figure 1B), thus selecting genetic variants in each individual that are predicted to affect both gene expression and TF binding. The effect of a SNP *s* on TF *i*’s regulation of gene *j* is then defined as the product 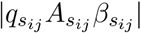. Modifier weights to the reference motif prior are calculated by adding these effects per TF-gene pair, allowing for the fact that a gene might have more than one variant in its promoter region affecting the binding of a particular TF. The EGRET prior network *E* is constructed by subtracting the modifier from the reference motif prior:

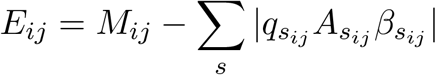

thus penalizing the reference motif prior when the individual in question contains a genetic variant with sufficient evidence to suggest it may alter gene regulation (Figure 1B, Supplementary Note S3). This EGRET prior, *E*, is then combined with two additional inputs (Supplementary Note S4). The fifth input to EGRET is a TF-TF interaction network (*P*) derived from protein-protein interactions, that reflects the fact that TFs can form complexes through protein-protein interactions to cooperatively regulate expression. Lastly, EGRET uses gene expression data to calculate a co-expression matrix *C* under the assumption that genes which are co-regulated are likely to exhibit correlated expression.

EGRET then uses a message passing network integration framework introduced previously [6] to search for regulatory consistency among the *E*, *P*, and *C* matrices (Supplementary Note S5). This message passing process updates all three input matrices, boosting those relationships that show agreement between associations captured in *E*, *P*, and *C* while down-weighting others. Upon convergence, the primary output is an individual-specific, complete, bipartite GRN (*E*\*) that captures genotype-specific regulatory effects. EGRET repeats this process separately using genotype information for each individual, producing a collection of individual-specific genotype-informed GRNs (Figure 1C). These networks can then be examined to identify features that are unique to specific genotypes, are associated with particular phenotypic states, or both. It is important to note that EGRET GRNs *E*\* are complete graphs, meaning that an edge exists between all TFs and genes considered. However, it is the edge weights that indicate the strength of the relationship between the respective TFs and genes, with a higher weight indicating a higher likelihood of a regulatory relationship.

It is also worth noting that the data for *M*, *P*, *C*, as well as the eQTLs can be obtained from publicly available resources. Thus, one can construct an EGRET network for a given cell type in an individual of interest simply by providing the genotype information for that individual and relying on publicly available data (for example, databases such as the Genotype Tissue Expression Project - GTEx [11]) for the remaining model inputs.

### 2.2 Regulatory disruption scores

EGRET inferred edge weights can be used to quantitatively estimate the predicted regulatory effects produced by SNPs on a given gene, TF, or TF-gene relationship (Table S2). A higher edge weight between a TF *i* and a gene *j* is interpreted as a higher confidence that the TF binds the promoter of and regulates the expression of gene *j*. To assess the effects of SNPs on gene regulation, we define and calculate three different regulatory disruption scores for nodes and edges in a given genotype *x* (Figure 1D). The *edge disruption score* 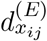 quantifies the extent to which a TF-gene regulatory relationship is disrupted by genetic variants. The *gene disruption score* 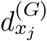 assesses the extent to which a gene has disrupted regulation due to genetic variants in its promoter region. The *TF disruption score* 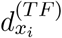 is a measure of the extent to which a TF’s genome-wide regulation is disrupted by genetic variants. These scores are defined per edge/node in each genotype-specific EGRET network by comparing it to a baseline network constructed using no genotype information and applying message-passing to *M*, *P*, and *C* (instead of *E*, *P*, and *C*):

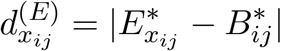

where 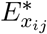 denotes the weight of edge *ij* in the EGRET network for individual *x* and 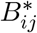 is the edge weight for edge *ij* in the baseline network predicted without using genotype information. This score quantifies the extent to which edges are disrupted by variants in a given individual-specific network (*E*\*) compared to a baseline genotype-agnostic regulatory network (*B*\*).

Similarly, TF disruption scores 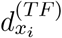 and gene disruption scores 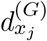 are calculated by taking the sum of edge disruption scores around the specific TF or gene in question:

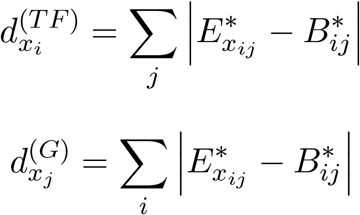

It is worth noting that disruption scores are all greater-than or equal to zero and that a higher edge disruption score corresponds to a larger difference between the EGRET and baseline edge weights for a particular TF-gene edge. However, because of the manner in which the EGRET prior is created - by penalizing edges involving a TF motif which contains an eQTL variant with a significant negative QBiC effect - only regulatory disruptions are modeled, and not the creation of “new” regulatory relationships where none potentially existed before.

## 3 Applications of EGRET

### 3.1 EGRET finds regulatory differences between two genetically distinct cell lines

We tested whether EGRET could distinguish genotype-specific patterns of gene regulation by analyzing two blood-derived cell lines, GM12878 and K562. We chose these cell lines because high quality genome sequences are available for both cell lines [12, 13] providing high confidence variant calls; this was especially useful for K562 because the cell line is aneuploid [13]. In addition, both cell lines have had relatively large numbers of TFs mapped by ChIP-seq (110 TFs for GM12878 and 204 TFs for K562 in the ReMap 2018 database [14]), allowing us to use differential TF binding as a way of validating regulatory differences.

To build genotype-specific EGRET priors (*E*) for GM12878 and K562, we generated a reference motif prior *M* using FIMO [9] identifying TF motifs in the promoter regions of genes ([−750, +250] relative to transcription start sites) and modified this using eQTL data for lymphoblastoid cell lines (LCLs) from GTEx [11], the cell lines’ respective genotypes, and SNP effect predictions from QBiC (Supplementary Notes S1-S3). Comparing edges in *E* against *M* predicted 1,520 genotype-altered prior edges for GM12878 and 1,182 for K562 (Figure S2) out of a total of 39,690,052 possible edges.

Next, we used the TF-TF interaction data as used by Sonawane and colleagues [15] to construct *P*, and LCL gene expression data from GTEx to construct *C* (Supplementary Note S4). Genes and TFs with low expression, defined as having non-zero values in < 50 samples, were filtered out. Performing message passing between *E*, *P*, and *C* produced the final genotype-specific EGRET networks *E*\* for GM12878 and K562 (Supplementary Note S5). For comparison, we constructed a baseline PANDA GRN using *M* as input to the message passing with *P* and *C*. We calculated the edge disruption score for each TF-gene pair in each cell line’s EGRET network. Because of the relatively small number of genotype-altered edges in the EGRET priors, the majority of edge disruption scores are very close to zero in both cell lines (Figure S3).

For each cell line, we compared both EGRET’s predictions of TF binding and the baseline PANDA networks to an empirical network based on ChIP-seq data [14] (Supplementary Note S6.1). At multiple cut-offs for the edge disruption scores, EGRET networks outperformed the baseline network prediction of TF binding for variant-disrupted edges (Tables S3 and S4, Supplementary Note S6.2). Based on these analyses, we considered variant-impacted scores to be those at or above 0.35 and a “high” disruption score to be anything at or above 0.5.

To capture changes in the disruption score between different genotype-specific networks, we calculated a “regulatory difference score” 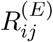 (Table S2) for each edge between genotypes GM12878 (*g*) and K562 (*k*), defined as:

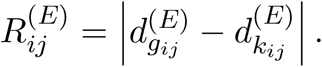

The magnitude of this score is the difference in edge disruption scores between GM12878 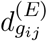 and K562 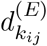 and reflects the assumption that genetic differences between cell lines will cause differences in predicted regulatory TF-gene interaction strength. A high value of 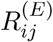 suggests a difference in the edge disruption scores for the edge *TF_i_ − G_j_* between the two cell lines, which is interpreted as that relationship being disrupted in one cell line but not the other. Conversely, a value of 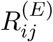 close to zero indicates that the edge disruption scores for the edge *TF_i_ − G_j_* are similar and therefore that the regulatory relationship is either disrupted in both cell lines or remains unaltered in both cell lines.

We again used the cell line-specific ChIP-seq regulatory networks (Supplementary Note S6.1) to construct a *differential ChIP-seq regulatory network* by taking the absolute value of the difference between the GM12878 ChIP-seq network and the K562 ChIP-seq network. This allows us to assign a score of 1 to edges which show differential TF binding (a TF binds the promoter region of a gene in one cell line but not the other) and a score of 0 to edges which show the same pattern of TF binding (a TF either binds the promoter region of a gene in both cell lines, or neither). This scoring allows us to validate the framework modeled by the regulatory difference score 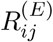. We found that edges with high differential regulation scores 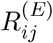 were enriched for edges showing differential TF binding in the differential ChIP-seq regulatory network (Fisher p-value = 2.4e-226, T-test p-value = 2.296e-07).

We highlight two examples of genotype-specific promoter binding of TFs identified though the EGRET network analysis. First, the edge between the TF RELA and the gene SLC16A9 (ENSG00000165449) has a regulatory difference score of 6.099744, with 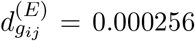 in GM12878 and 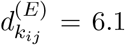 in the K562. These scores suggest that the binding of RELA to the promoter region of SLC16A9 is disrupted in K562, but not in GM12878. The positions of eQTLs, genetic variants, and ChIP-seq binding regions for RELA in both genotypes (Figure 2A) indicate that an eQTL variant is present in the promoter region of SLC16A9 (purple track in Figure 2A), is associated with the expression of SLC16A9, resides within a RELA binding motif, and is predicted by QBiC to affect the binding of RELA at that location; the disrupting variant is present only in K562 (orange track in Figure 2A) and not in GM12878; this prediction is confirmed by the presence of a RELA ChIP-seq binding range in GM12878, but not in K562 (teal track in Figure 2A). As a second example, consider the edge between of the TF ARID3A and the gene PMS2CL (ENSG00000187953), with a regulatory difference score of 1.0564 and 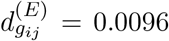 in GM12878 and 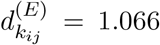 in K562, suggesting that the binding of ARID3A to the promoter region of PMS2CL is disrupted in K562, but not in GM12878. This prediction is confirmed by ChIP-seq-derived TF binding data in the region (Figure 2B). Both of these examples of genotype-specific TF binding are within the top 20 edge disruption scores for K562 for edges with confirmed differential binding between K562 and GM12878 ChIP-seq experiments.

**Figure 2:**
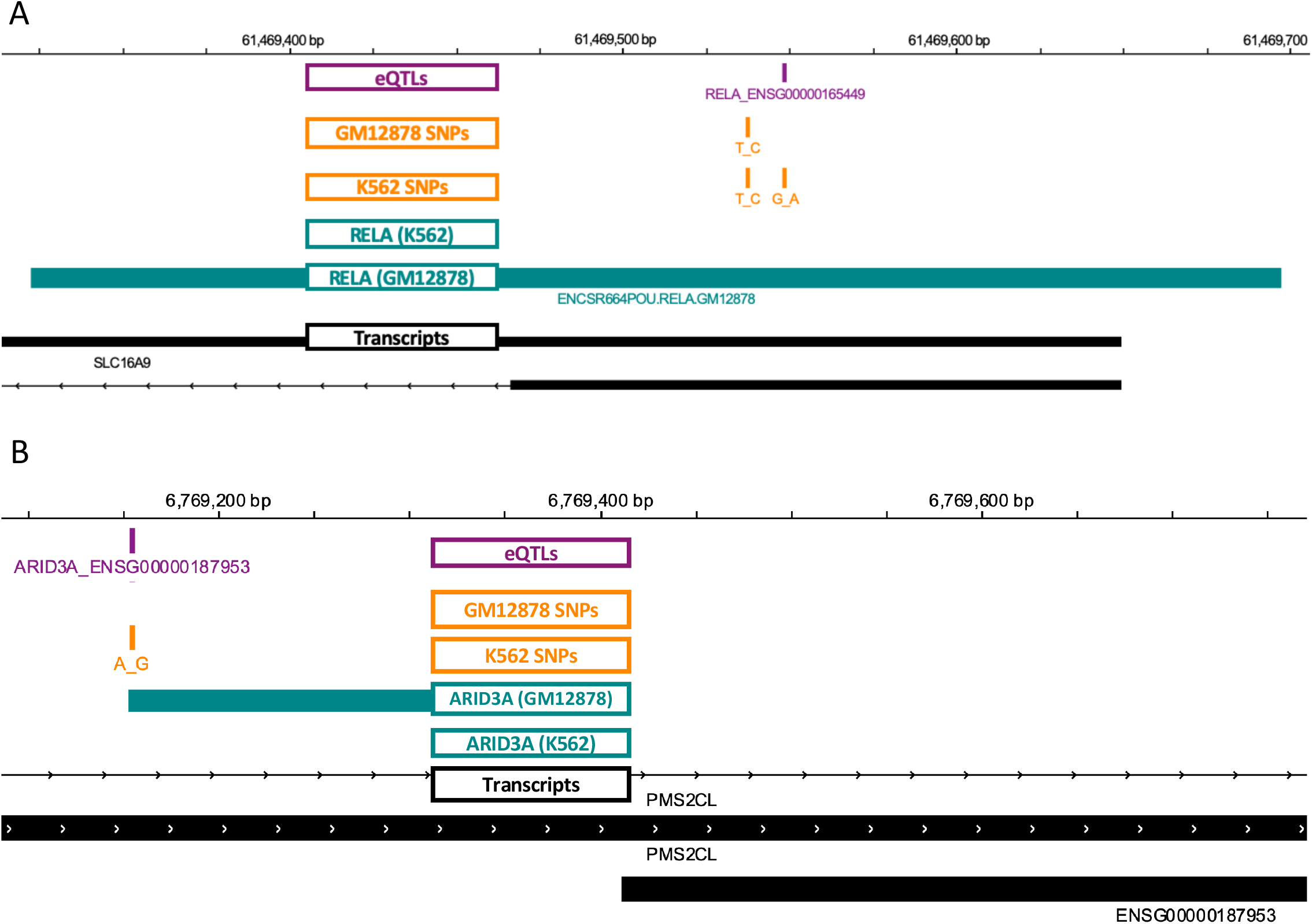
EGRET identifies variant-impacted TF binding disruptions. **(A)** Example of variant-disrupted RELA binding in K562 but not in GM12878. Positions of eQTLs (purple track), genetic variants (orange tracks), ChIP-seq binding regions (teal tracks), and genes (black track) are shown in the region of SLC16A9. **(B)** Example of variant-disrupted ARID3A binding in K562 but not in GM12878. Positions of eQTLs (purple track), genetic variants (orange tracks), ChIP-seq binding regions (teal tracks) and genes (black track) are shown in the region of PMS2CL. The eQTL track is labeled according to the TF motif in which the eSNP resides as well as the adjacent eGene.

Because an edge in an EGRET network implies a *regulatory* relationship, as opposed to simply presence of binding by a TF, we wanted to further validate our network predictions against assays that captured changes in gene expression. We used data from an in-vitro allele-specific expression (ASE) assay (Biallelic Targeted Self-Transcribing Active Regulatory Region sequencing — BiT-STARR-seq) performed in LCLs [16] (Supplementary Note S6.3). We calculated regulatory difference scores per gene and found that the 101 genes having highest differential regulation scores 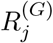 (those within the top 10%) were enriched for genes harboring ASE-causing variants located within promoter region TF motifs (Fisher p-value = 2.5e-03). As a second independent validation, we compared data from a published chromatin accessibility QTL (caQTL) analysis in LCLs to those genes whose regulatory difference score were in the top 10% (Supplementary Note S6.4) [17] and found that these genes having 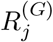 values among the top 10% were enriched for having caQTLs within motifs in their promoter regions (Fisher p-value = 1.4e-04). This suggests that many of the predicted regulatory SNPs alter their associated regulatory networks by affecting chromatin accessibility. It is worth noting that our results are based only on the genotypes of two cell lines; we anticipate that using a larger number of genotyped cell lines with available ChIP-seq, caQTL, and ASE data would increase both the specificity and sensitivity of predicting genotype-mediated effects in gene expression.

Overall, these results indicate that EGRET is capable of synthesizing diverse sources of data to model gene regulatory processes and can predict genotype-associated patterns of gene regulation.

### 3.2 EGRET networks for a population of individuals identify cell type-specific disease associations

A growing body of work indicates that cell-type specific gene regulatory processes affect gene expression [18, 15] and do so in a manner dependent on an individual’s genotype [19, 20, 21], resulting in changes that alter the structure of functional “communities” or “modules” comprised of TFs and genes, and are enriched for genes associated with tissue-specific biological processes [22]. Banovich and colleagues [17] had previously analyzed RNA-seq data derived from three cell types: lymphoblastoid cell lines (LCLs), induced pluripotent stem cells (iPSCs), and cardiomyocytes (CMs; differentiated from the iPSCs). They demonstrated that genes preferentially expressed in CMs were enriched for processes associated with coronary artery disease, and those enriched in LCLs were associated with immune-related conditions. Our working hypothesis was that these effects should be linked to cell type-specific regulatory processes affected by an individual’s genetic background.

To test this, we constructed 357 individual-specific EGRET networks using expression, genotype, and eQTL data from 119 Yoruba individuals for all three cell types used in the Banovich *et al.* [17] study (Supplementary Note S7). We also constructed a baseline GRN for each cell type (Supplementary Note S7.1). We calculated TF disruption scores (defined in Supplementary Table S2) for each TF in each individual EGRET network to identify TFs whose regulatory influence was disrupted by variants. TF disruption scores 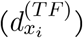 were then scaled per individual and cell type to have a mean of zero and standard deviation of one, and are denoted 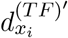 (Supplementary Note S7.2). We then labelled TFs as associated with Crohn’s disease (CD) and coronary artery disease (CAD) (Tables S5 and S6) based on annotation from the NHGRI-EBI GWAS catalog [23]. We tested to see if disease-associated TFs were more likely to have significant disruption scores in relevant cell types. Using a T-test, we found that TF disruption scores were significantly higher in cardiomyocytes (CMs) for TFs associated with CAD than were disruption scores for non-CAD related TFs (p = 4.5256e-06); this CAD enrichment was not observed in LCLs (p = 0.99831). Similarly, we found TF disruption scores in LCLs, but not CMs, were substantially higher for TFs linked to CD than for non CD-linked TFs (p = 5.3374e-16 in LCL networks, p = 1 in CM networks; Table 2). This analysis leads to an important observation: genotype-mediated, disease-related TF disruptions are cell-type specific and can be identified using networks inferred by EGRET. Indeed, we find that the highest TF disruption scores for CAD TFs occur in CMs (Figure 3A) and that the highest TF disruption scores for CD TFs occur in LCLs (Figure 3B).

**Table 2:** T-test p-values of differences between the TF disruption scores of disease (CD or CAD related TFs, determined from the GWAS catalog) versus non-disease TFs in different cell types. CAD TFs have significantly higher TF disruption scores than non-CAD TFs in CMs, but not in LCLs. CD TFs have significantly higher TF disruption scores than non-CD TFs in LCLs, but not in CMs.

| Disease | Cell type | P-value |
| --- | --- | --- |
| CAD | LCL | 0.99831 |
| CAD | CM | 4.5256e-06 |
| CD | LCL | 5.3374e-16 |
| CD | CM | 1 |

**Figure 3:**
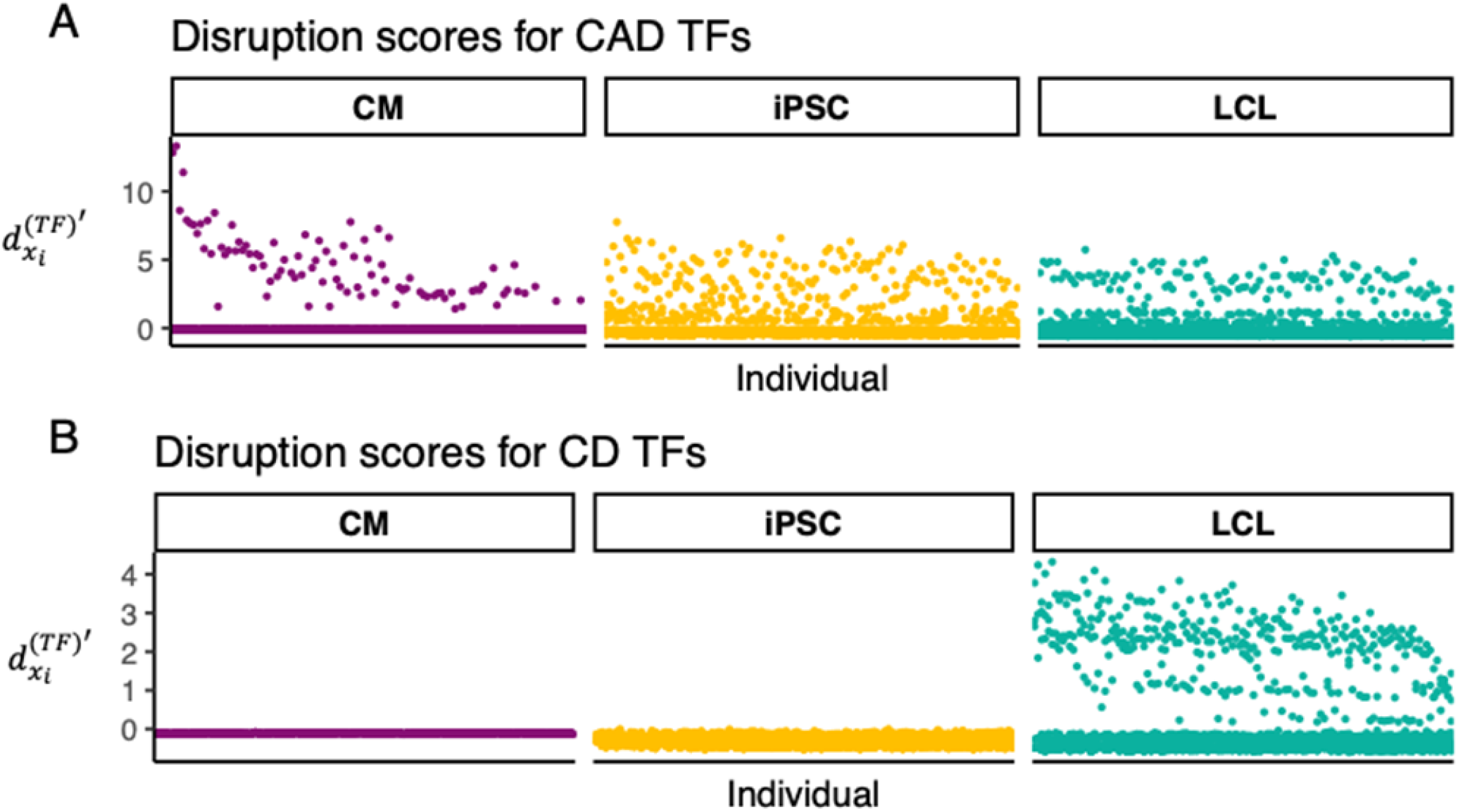
Disease-related TFs are disrupted in relevant cell types. Scaled TF disruption scores 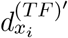 are shown for 119 Yoruba individuals for TFs associated with **(A)** coronary artery disease (CAD) or **(B)** Crohn’s disease (CD). Each point represents the scaled TF disruption score for a disease-related (CD or CAD) TF, for a given individual for a given cell type (LCL, CM or iPSC). Disease-related TFs were identified using the GWAS catalog [23]. Scaled TF disruption scores for CAD-related TFs are highest in the cardiac-related cell type, CMs. Scaled TF disruption scores for CD-related TFs are highest in the immune cell type, LCLs.

Further supporting this observation, the TF disruption signal in CAD is dominated in a subset of the study population by single a TF, ERG, which is a member of the erythroblast transformation-specific (ETS) gene family and known to be involved in angiogenesis [24]. In these individuals, the high TF disruption scores for CAD TFs in CMs are driven by the presence or absence of a mutation on chromosome 1 (chr1:201476815, an eQTL for CSRP1) that lies in the binding motif for the TF ERG in the promoter region of the gene CSRP1 (ENSG00000159176). While ERG is identified as CAD-related in the GWAS catalog, CSRP1 (alias CRP1) is not. However, CSRP1 is a known smooth muscle marker [25] and has been found by GTEx [11] to be highly expressed in smooth muscles, especially in arteries (Figure S4). CSRP1 has also been associated with the bundling of actin filaments [26], cardiovascular development [27], and with response to arterial injury [28]. Further, knockdown of CSRP1 in zebrafish caused cardiac bifida [29] and a frameshift mutation in CSRP1 has been linked to congenital cardiac defects in a large human pedigree [30]. The results of our EGRET analysis support a previously unreported mechanism of action for ERG in heart disease—that ERG regulates the expression of CSRP1 and that this regulation can be disrupted by genetic variation.

We also tested the hypothesis that the network effects of genetic variants have the potential to subtly change the modular structure of genotype-specific networks, altering the functional network modules active in an individual. ALPACA [22] is a method that compares the modular structure of two networks and identifies modules that differ between the networks. The resulting gene differential modularity (DM) scores indicate which genes have undergone the greatest change in their “modular environment.” We used ALPACA to compare the modular structure of the cell-type and individual-specific EGRET GRNs with the baseline GRN for the corresponding cell type, and calculated the DM score for each gene in each network (Supplementary Note S7.3, Figure S5).

Given that individual 18 had the greatest TF disruption score for ERG in CMs, we further investigated cellular processes predicted by EGRET to be variant-perturbed within this individual’s three cell-type specific EGRET networks. For each cell type we ranked this individual’s genes by their DM scores from highest to lowest in each cell type reflecting their predicted impact on altering the modular structure of each cell-type specific network. We used GORILLA (Supplementary Note S7.4; [31]) with these ranked lists to identify GO biological process functions associated with modules altered by the presence of specific genetic variants. Several GO terms relevant to CMs and cardiovascular functioning and development, including “regulation of actomyosin structure organization,” “prepulse inhibition,” “ephrin receptor signaling pathway,” “maintenance of postsynaptic specialization structure,” and “actin cytoskeleton reorganization” were enriched in CMs from this individual (Figure 4, Table S7) but not in their LCLs or iPSCs (Figure 4, Tables S8 and S9). For full enrichment results, see Figure S6, Figure S7, and S8, Figure S9). Further evidence of cell type-specific alteration of functional modules can be seen by examining the DM scores of disease-associated target genes (as annotated by the NHGRI-EBI GWAS catalog [23]). Coronary artery disease genes with high DM scores in CMs had low DM scores in iPSCs and LCLs (Figure 5A). In contrast, genes associated with Crohn’s disease, which has a strong immune component, that had high DM scores in LCLs and low DM scores in iPSCs and CMs (Figure 5B).

**Figure 4:**
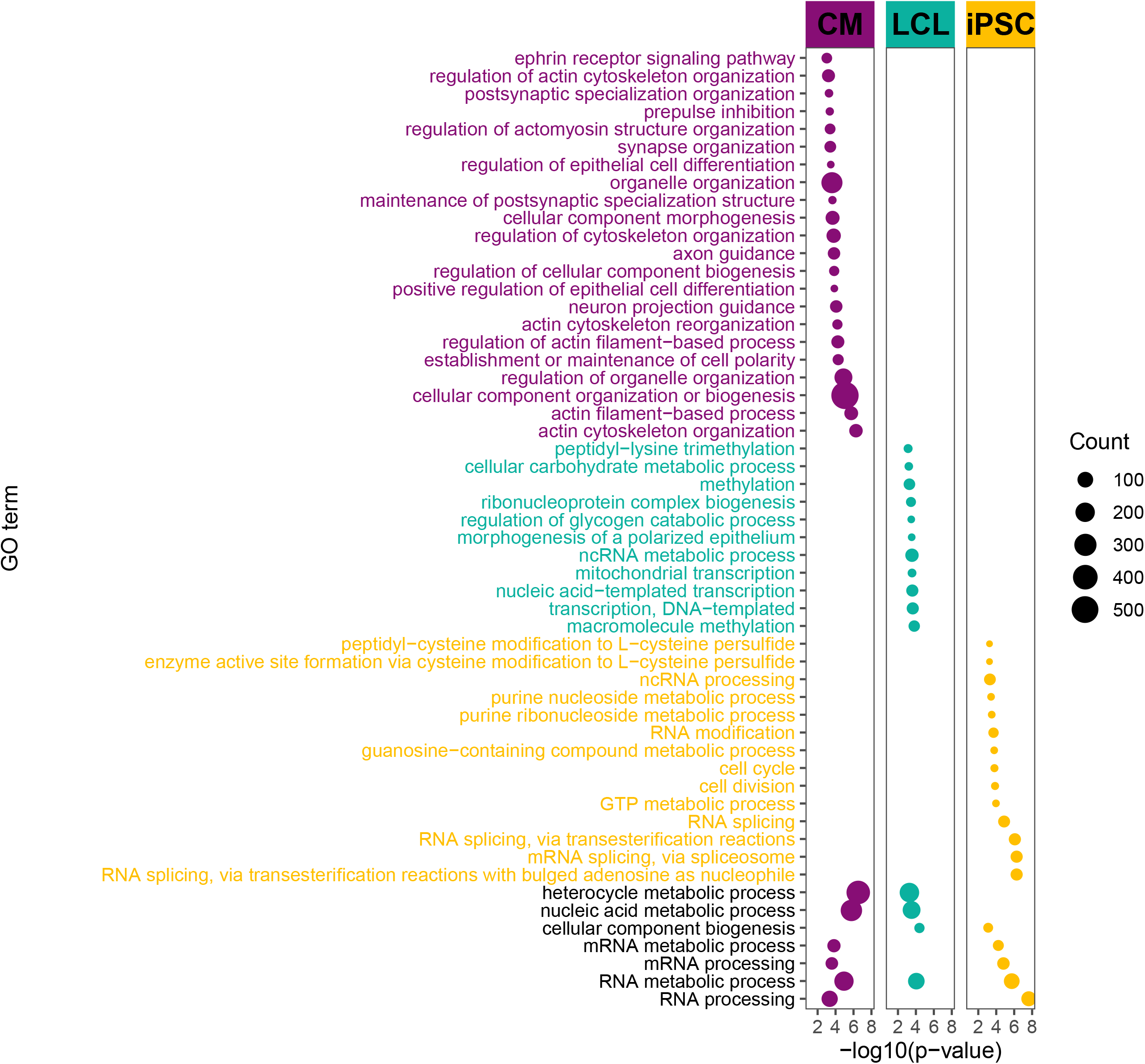
Variant-disrupted gene regulation affecting network modularity is enriched for coronary/heart related functions in CMs for an individual with a CAD disruption signature. GO terms enriched in genes with high DM scores for individual 18, the individual with the highest TF disruption score for ERG. Several GO terms related to coronary/cardiac function are enriched in highly ranked DM genes in CMs but not in LCLs and iPSCs. Point size corresponds to the the number of high-DM genes annotated with the corresponding GO term. For display purposes, several generic GO terms enriched only in CMs were omitted in this figure. The entire set of enriched GO terms can be seen in Figure S9, as well as Tables S7, S9 and S8

**Figure 5:**
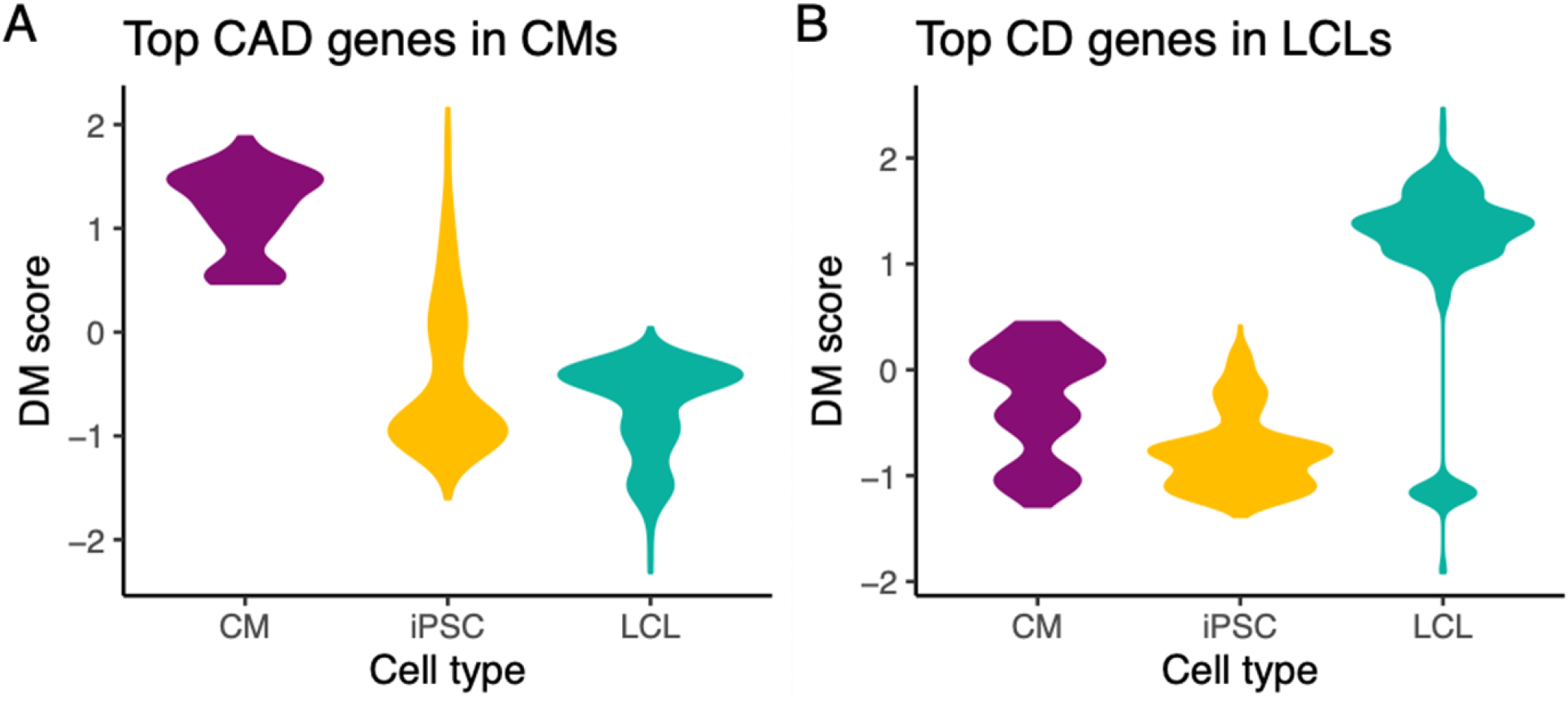
Variant-disrupted disease genes affect the modularity of the individual’s regulatory network in the relevant cell type. Differential modularity (DM) scores indicate the extent to which a gene’s modular environment in the network changes between the genotype-specific EGRET network and the genotype-agnostic network. **(A)** CAD-related genes with high DM scores in cardiomyocytes (CMs) have low DM scores in the other cell types; **(B)** CD-related genes with high DM scores in LCLs have low scores in the other cell types.

EGRET also predicts dosage effects of regulatory SNP variants on network structure. Consider CSRP1, which we previously discussed as having a regulatory SNP in its promoter region that can affect binding of the transcription factor ERG. EGRET shows that the presence of a genetic variant in the promoter region of CSRP1 affects not only regulation by ERG (as seen by a substantial TF disruption score) but also the role that CSRP1 plays in altering the functional modules in cardiomyocyte GRN models. As seen by CSRP1’s DM scores in Figure S10, EGRET predicts that the genetic variant exerts its influence on network structure in a dosage-specific manner; individuals homozygous for the disrupting variant are predicted to exhibit the greatest impact on the modularity, those who are heterozygous to have an intermediate effect, and those homozygous for the wild-type to exhibit minimal or no effect on modularity.

Collectively, these results suggest that phenotype- and disease-associated variants can act through disruption of TF binding leading to regulatory changes that manifest themselves both through altered expression of specific target genes and the modification of GRN functional modular structure.

## 4 Discussion

One of the fundamental tenets of genetics is that genotype influences phenotype. For many traits, especially those related to human disease, this connection is not straightforward. The vast majority of phenotype-associated genetic variants are non-coding and have small effect sizes [32, 33] and a recent analysis found that most (71%-100%) 1-MB windows in the genome contribute to schizophrenia heritability [34]. This suggests that many variants must act in concert to produce complex trait phenotypes, but the mechanisms by which they exert their influence remains an open question. Functional genomics studies have provided some insights into roles of these variants: variants are enriched in regulatory elements [35, 36, 37], disease heritability tends to be enriched in tissues relevant to the disease [1], and TF binding plays an important role in explaining heritability of human traits [5]. Despite this progress at the population level, questions remain regarding the influence of an individual’s genotype on these regulatory processes. Answers to these questions will be important for translating population-level insights into clinically actionable information.

EGRET is the first method, to our knowledge, that directly addresses these issues. EGRET begins with a reference motif prior network based on mapping transcription factor binding sites to the regulatory regions of genes. EGRET extends this by modifying the reference motif network based on evidence that SNPs in a gene’s regulatory region may influence TF binding as well as gene expression. Subsequently, EGRET uses a previously developed message passing framework [6] to iteratively seek consistency between a genotype-altered regulatory network model, TF-TF protein-protein interaction data (acknowledging that TFs can form regulatory complexes), and gene co-expression data (based on the assumption that genes regulated by the same TFs are likely to exhibit correlated expression). EGRET then outputs edge weights for all TF-to-gene edges. These edge weights reflect the confidence of a regulatory relationship between a TF and gene, given an individual’s genotype. We demonstrate EGRET using publicly available eQTL, gene expression, and PPI data, and show that the algorithm provides powerful insights regardless of whether or not the individual genotypes are sample matched with other data types.

We validated EGRET in two ways. In the first, we inferred genotype-specific gene regulatory networks for two genotyped cell lines and identified genes that differed in their TF-gene edges, meaning that the model predicts differences in binding of specific TFs to upstream regions of individual genes. When we cross-referenced EGRET’s predictions with ChIP-seq data for these cell lines, we found concordance between the predictions and ChIP-seq data, demonstrating that EGRET was able to accurately identify different TF binding patterns and effectively altered the structure of the regulatory network. We also found that genes with high regulatory difference scores between the two cell lines—those predicted to be differently regulated by EGRET—were enriched for QTLs associated with chromatin accessibility and enriched for allele-specific expression, suggesting that the EGRET-predicted regulatory changes are likely to have broader regulatory effects.

Our second validation looked at three different cell types in 119 genotyped individuals. We found distinct cell type-specific and genotype-specific differences in the gene regulatory networks that were linked to disease. Most notable among these were regulatory differences associated with Crohn’s disease in lymphoblastoid cell lines and others linked to coronary artery disease in the regulatory networks in cardiomyocytes. Not only were individual TF-gene connections disrupted, but these disruptions led to higher-order changes in the network community structure, reorganizing the network in ways that predict changes in cell type- and disease-specific functional network communities.

Taken together, these results from EGRET present a compelling picture of the way in which small-effect, non-coding SNPs work together to influence phenotype. These SNPs have the potential to subtly alter the binding of TFs to their target genes. The direct effect of these individual SNPs is to alter which TFs regulate specific genes. However, their indirect, and possibly more important effect, is to alter the structure and membership of functional communities in the overall regulatory networks. Indeed, it is known that even a small number of TF-gene regulatory edge additions or deletions can lead to significant changes in network modular organization [22].

EGRET is capable of inferring gene regulatory networks specific to an individual’s genotype, synthesizing genetic and gene expression data in a way that, for the first time, allows verifiable, disease-associated regulatory changes to be inferred for individual research subjects. As such, EGRET has the potential to substantially advance our understanding of genetic effects on disease risk, development, and response to therapeutic interventions. Potential applications of EGRET are wide ranging. EGRET can be used to infer a specific gene regulatory network for any individual for whom genotype data are available, even without associated gene expression data—provided there is expression and eQTL data from a relevant cell type obtained from a sufficiently large population to infer accurate regulatory network models. This implies that EGRET can be used to retrospectively analyze large cohort GWAS studies to tease out mechanistic associations for phenotype-linked genetic variants, as well as in the context of new studies that seek to understand disease mechanisms and the regulatory role of non-coding genetic variants.

## Data and Code Availability

EGRET is available through the Network Zoo R package (netZooR v0.9; netzoo.github.io) with a step-by-step tutorial.

## 5 Funding

DW, MBG, and JQ are supported through a grant from the US National Cancer Institute, 1R35CA220523. MBG and JQ also supported by 1U24CA231846. JP is supported by a grant from the US National Heart, Lung and Blood Institute, K25HL140186. KG is supported by a grant from the US National Heart, Lung and Blood Institute, K25HL133599.

## 6 Author Contributions

DW and JP developed the EGRET method, DW implemented, tested and applied the method, DW and JP interpreted the results, MBG assisted with software distribution and manuscript preparation, JQ and KG provided input into all stages of the project, JP conceived of the study.

## Notes

https://netzoo.github.io/

### Competing Interest Statement

The authors have declared no competing interest.

## References

[1] Evan A. Boyle, Yang I. Li, and Jonathan K. Pritchard. An expanded view of complex traits: From polygenic to omnigenic. Cell, 169(7):1177 – 1186, 2017.

[2] Alexander Gusev, Nicholas Mancuso, Hyejung Won, Maria Kousi, Hilary K. Finucane, Yakir Reshef, Lingyun Song, Alexias Safi, Steven McCarroll, Benjamin M. Neale, Roel A. Ophoff, Michael C. O’Donovan, Gregory E. Crawford, Daniel H. Geschwind, Nicholas Katsanis, Patrick F. Sullivan, Bogdan Pasaniuc, Alkes L. Price, and Schizophrenia Working Group of the Psychiatric Genomics Consortium. Transcriptome-wide association study of schizophrenia and chromatin activity yields mechanistic disease insights. Nature Genetics, 50(4):538–548, 2018.

[3] Hilary K Finucane, Brendan Bulik-Sullivan, Alexander Gusev, Gosia Trynka, Yakir Reshef, Po-Ru Loh, Verneri Anttila, Han Xu, Chongzhi Zang, Kyle Farh, Stephan Ripke, Felix R Day, Shaun Purcell, Eli Stahl, Sara Lindstrom, John R B Perry, Yukinori Okada, Soumya Raychaudhuri, Mark J Daly, Nick Patterson, Benjamin M Neale, Alkes L Price, ReproGen Consortium, Schizophrenia Working Group of the Psychiatric Genomics Consortium, and The RACI Consortium. Partitioning heritability by functional annotation using genome-wide association summary statistics. Nature Genetics, 47(11):1228–1235, 2015.

[4] Zhihong Zhu, Futao Zhang, Han Hu, Andrew Bakshi, Matthew R Robinson, Joseph E Powell, Grant W Montgomery, Michael E Goddard, Naomi R Wray, Peter M Visscher, and Jian Yang. Integration of summary data from gwas and eqtl studies predicts complex trait gene targets. Nat Genet, 48(5):481–487, May 2016.

[5] Bryce van de Geijn, Hilary Finucane, Steven Gazal, Farhad Hormozdiari, Tiffany Amariuta, Xuanyao Liu, Alexander Gusev, Po-Ru Loh, Yakir Reshef, Gleb Kichaev, Soumya Raychauduri, and Alkes L Price. Annotations capturing cell type-specific TF binding explain a large fraction of disease heritability. Human Molecular Genetics, 29(7):1057–1067, 10 2019.

[6] Kimberly Glass, Curtis Huttenhower, John Quackenbush, and Guo-Cheng Yuan. Passing messages between biological networks to refine predicted interactions. PloS one, 8(5):e64832, 2013.

[7] Daniel Taliun, Daniel N. Harris, Michael D. Kessler, Jedidiah Carlson, Zachary A. Szpiech, Raul Torres, Sarah A. Gagliano Taliun, André Corvelo, Stephanie M. Gogarten, Hyun Min Kang, Achilleas N. Pitsillides, Jonathon LeFaive, Seung-been Lee, Xiaowen Tian, Brian L. Browning, Sayantan Das, Anne-Katrin Emde, Wayne E. Clarke, Douglas P. Loesch, Amol C. Shetty, Thomas W. Blackwell, Albert V. Smith, Quenna Wong, Xiaoming Liu, Matthew P. Conomos, Dean M. Bobo, François Aguet, Christine Albert, Alvaro Alonso, Kristin G. Ardlie, Dan E. Arking, Stella Aslibekyan, Paul L. Auer, John Barnard, R. Graham Barr, Lucas Barwick, Lewis C. Becker, Rebecca L. Beer, Emelia J. Benjamin, Lawrence F. Bielak, John Blangero, Michael Boehnke, Donald W. Bowden, Jennifer A. Brody, Esteban G. Burchard, Brian E. Cade, James F. Casella, Brandon Chalazan, Daniel I. Chasman, Yii-Der Ida Chen, Michael H. Cho, Seung Hoan Choi, Mina K. Chung, Clary B. Clish, Adolfo Correa, Joanne E. Curran, Brian Custer, Dawood Darbar, Michelle Daya, Mariza de Andrade, Dawn L. DeMeo, Susan K. Dutcher, Patrick T. Ellinor, Leslie S. Emery, Celeste Eng, Diane Fatkin, Tasha Fingerlin, Lukas Forer, Myriam Fornage, Nora Franceschini, Christian Fuchsberger, Stephanie M. Fullerton, Soren Germer, Mark T. Gladwin, Daniel J. Gottlieb, Xiuqing Guo, Michael E. Hall, Jiang He, Nancy L. Heard-Costa, Susan R. Heckbert, Marguerite R. Irvin, Jill M. Johnsen, Andrew D. Johnson, Robert Kaplan, Sharon L. R. Kardia, Tanika Kelly, Shannon Kelly, Eimear E. Kenny, Douglas P. Kiel, Robert Klemmer, Barbara A. Konkle, Charles Kooperberg, Anna Köttgen, Leslie A. Lange, Jessica Lasky-Su, Daniel Levy, Xihong Lin, Keng-Han Lin, Chunyu Liu, Ruth J. F. Loos, Lori Garman, Robert Gerszten, Steven A. Lubitz, Kathryn L. Lunetta, Angel C. Y. Mak, Ani Manichaikul, Alisa K. Manning, Rasika A. Mathias, David D. McManus, Stephen T. McGarvey, James B. Meigs, Deborah A. Meyers, Julie L. Mikulla, Mollie A. Minear, Braxton D. Mitchell, Sanghamitra Mohanty, May E. Montasser, Courtney Montgomery, Alanna C. Morrison, Joanne M. Murabito, Andrea Natale, Pradeep Natarajan, Sarah C. Nelson, Kari E. North, Jeffrey R. O’Connell, Nicholette D. Palmer, Nathan Pankratz, Gina M. Peloso, Patricia A. Peyser, Jacob Pleiness, Wendy S. Post, Bruce M. Psaty, D. C. Rao, Susan Redline, Alexander P. Reiner, Dan Roden, Jerome I. Rotter, Ingo Ruczinski, Chloé Sarnowski, Sebastian Schoenherr, David A. Schwartz, Jeong-Sun Seo, Sudha Seshadri, Vivien A. Sheehan, Wayne H. Sheu, M. Benjamin Shoemaker, Nicholas L. Smith, Jennifer A. Smith, Nona Sotoodehnia, Adrienne M. Stilp, Weihong Tang, Kent D. Taylor, Marilyn Telen, Timothy A. Thornton, Russell P. Tracy, David J. Van Den Berg, Ramachandran S. Vasan, Karine A. Viaud-Martinez, Scott Vrieze, Daniel E. Weeks, Bruce S. Weir, Scott T. Weiss, Lu-Chen Weng, Cristen J. Willer, Yingze Zhang, Xutong Zhao, Donna K. Arnett, Allison E. Ashley-Koch, Kathleen C. Barnes, Eric Boerwinkle, Stacey Gabriel, Richard Gibbs, Kenneth M. Rice, Stephen S. Rich, Edwin K. Silverman, Pankaj Qasba, Weiniu Gan, Namiko Abe, Laura Almasy, Seth Ament, Peter Anderson, Pramod Anugu, Deborah Applebaum-Bowden, Tim Assimes, Dimitrios Avramopoulos, Emily Barron-Casella, Terri Beaty, Gerald Beck, Diane Becker, Amber Beitelshees, Takis Benos, Marcos Bezerra, Joshua Bis, Russell Bowler, Ulrich Broeckel, Jai Broome, Karen Bunting, Carlos Bustamante, Erin Buth, Jonathan Cardwell, Vincent Carey, Cara Carty, Richard Casaburi, Peter Castaldi, Mark Chaffin, Christy Chang, Yi-Cheng Chang, Sameer Chavan, Bo-Juen Chen, Wei-Min Chen, Lee-Ming Chuang, Ren-Hua Chung, Suzy Comhair, Elaine Cornell, Carolyn Crandall, James Crapo, Jeffrey Curtis, Coleen Damcott, Sean David, Colleen Davis, Lisa de las Fuentes, Michael DeBaun, Ranjan Deka, Scott Devine, Qing Duan, Ravi Duggirala, Jon Peter Durda, Charles Eaton, Lynette Ekunwe, Adel El Boueiz, Serpil Erzurum, Charles Farber, Matthew Flickinger, Chris Frazar, Mao Fu, Lucinda Fulton, Shanshan Gao, Yan Gao, Margery Gass, Bruce Gelb, Xiaoqi Priscilla Geng, Mark Geraci, Auyon Ghosh, Chris Gignoux, David Glahn, Da-Wei Gong, Harald Goring, Sharon Graw, Daniel Grine, C. Charles Gu, Yue Guan, Namrata Gupta, Jeff Haessler, Nicola L. Hawley, Ben Heavner, David Herrington, Craig Hersh, Bertha Hidalgo, James Hixson, Brian Hobbs, John Hokanson, Elliott Hong, Karin Hoth, Chao Agnes Hsiung, Yi-Jen Hung, Haley Huston, Chii Min Hwu, Rebecca Jackson, Deepti Jain, Min A. Jhun, Craig Johnson, Rich Johnston, Kimberly Jones, Sekar Kathiresan, Alyna Khan, Wonji Kim, Greg Kinney, Holly Kramer, Christoph Lange, Ethan Lange, Leslie Lange, Cecelia Laurie, Meryl LeBoff, Jiwon Lee, Seunggeun Shawn Lee, Wen-Jane Lee, David Levine, Joshua Lewis, Xiaohui Li, Yun Li, Henry Lin, Honghuang Lin, Keng Han Lin, Simin Liu, Yongmei Liu, Yu Liu, James Luo, Michael Mahaney, and NHLBI Trans-Omics for Precision Medicine (TOPMed) Consortium. Sequencing of 53,831 diverse genomes from the nhlbi topmed program. Nature, 590(7845):290–299, 2021.

[8] Clare Bycroft, Colin Freeman, Desislava Petkova, Gavin Band, Lloyd T Elliott, Kevin Sharp, Allan Motyer, Damjan Vukcevic, Olivier Delaneau, Jared O’Connell, et al. The uk biobank resource with deep phenotyping and genomic data. Nature, 562(7726):203–209, 2018.

[9] Charles E Grant, Timothy L Bailey, and William Stafford Noble. Fimo: scanning for occurrences of a given motif. Bioinformatics, 27(7):1017–1018, 2011.

[10] Vincentius Martin, Jingkang Zhao, Ariel Afek, Zachery Mielko, and Raluca Gordân. Qbic-pred: quantitative predictions of transcription factor binding changes due to sequence variants. Nucleic acids research, 47(W1):W127–W135, 2019.

[11] John Lonsdale, Jeffrey Thomas, Mike Salvatore, Rebecca Phillips, Edmund Lo, Saboor Shad, Richard Hasz, Gary Walters, Fernando Garcia, Nancy Young, et al. The genotype-tissue expression (GTEx) project. Nature genetics, 45(6):580, 2013.

[12] Michael A Eberle, Epameinondas Fritzilas, Peter Krusche, Morten Källberg, Benjamin L Moore, Mitchell A Bekritsky, Zamin Iqbal, Han-Yu Chuang, Sean J Humphray, Aaron L Halpern, et al. A reference data set of 5.4 million phased human variants validated by genetic inheritance from sequencing a three-generation 17-member pedigree. Genome research, 27(1):157–164, 2017.

[13] Bo Zhou, Steve S Ho, Stephanie U Greer, Xiaowei Zhu, John M Bell, Joseph G Arthur, Noah Spies, Xianglong Zhang, Seunggyu Byeon, Reenal Pattni, et al. Comprehensive, integrated, and phased whole-genome analysis of the primary encode cell line k562. Genome research, 29(3):472–484, 2019.

[14] Jeanne Chèneby, Marius Gheorghe, Marie Artufel, Anthony Mathelier, and Benoit Ballester. Remap 2018: an updated atlas of regulatory regions from an integrative analysis of dna-binding chip-seq experiments. Nucleic acids research, 46(D1):D267–D275, 2018.

[15] Abhijeet Rajendra Sonawane, John Platig, Maud Fagny, Cho-Yi Chen, Joseph Nathaniel Paulson, Camila Miranda Lopes-Ramos, Dawn Lisa DeMeo, John Quackenbush, Kimberly Glass, and Marieke Lydia Kuijjer. Understanding tissue-specific gene regulation. Cell reports, 21(4):1077–1088, 2017.

[16] Cynthia A Kalita, Christopher D Brown, Andrew Freiman, Jenna Isherwood, Xiaoquan Wen, Roger Pique-Regi, and Francesca Luca. High-throughput characterization of genetic effects on dna–protein binding and gene transcription. Genome research, 28(11):1701–1708, 2018.

[17] Nicholas E Banovich, Yang I Li, Anil Raj, Michelle C Ward, Peyton Greenside, Diego Calderon, Po Yuan Tung, Jonathan E Burnett, Marsha Myrthil, Samantha M Thomas, et al. Impact of regulatory variation across human ipscs and differentiated cells. Genome research, 28(1):122–131, 2018.

[18] Camila M Lopes-Ramos, Marieke L Kuijjer, Shuji Ogino, Charles S Fuchs, Dawn L DeMeo, Kimberly Glass, and John Quackenbush. Gene regulatory network analysis identifies sex-linked differences in colon cancer drug metabolism. Cancer research, 78(19):5538–5547, 2018.

[19] Maud Fagny, Joseph N Paulson, Marieke L Kuijjer, Abhijeet R Sonawane, Cho-Yi Chen, Camila M Lopes-Ramos, Kimberly Glass, John Quackenbush, and John Platig. Exploring regulation in tissues with eqtl networks. Proceedings of the National Academy of Sciences, 114(37):E7841–E7850, 2017.

[20] GTEx Consortium et al. The gtex consortium atlas of genetic regulatory effects across human tissues. Science, 369(6509):1318–1330, 2020.

[21] Sarah Kim-Hellmuth, François Aguet, Meritxell Oliva, Manuel Muñoz-Aguirre, Silva Kasela, Valentin Wucher, Stephane E Castel, Andrew R Hamel, Ana Viñuela, Amy L Roberts, et al. Cell type–specific genetic regulation of gene expression across human tissues. Science, 369(6509), 2020.

[22] Megha Padi and John Quackenbush. Detecting phenotype-driven transitions in regulatory network structure. NPJ systems biology and applications, 4(1):1–12, 2018.

[23] Annalisa Buniello, Jacqueline A L MacArthur, Maria Cerezo, Laura W Harris, James Hayhurst, Cinzia Malangone, Aoife McMahon, Joannella Morales, Edward Mountjoy, Elliot Sollis, et al. The nhgri-ebi gwas catalog of published genome-wide association studies, targeted arrays and summary statistics 2019. Nucleic acids research, 47(D1):D1005–D1012, 2019.

[24] Aarti V Shah, Graeme M Birdsey, and Anna M Randi. Regulation of endothelial homeostasis, vascular development and angiogenesis by the transcription factor erg. Vascular pharmacology, 86:3–13, 2016.

[25] James R Henderson, Teresita Macalma, Doris Brown, James A Richardson, Eric N Olson, and Mary C Beckerle. The lim protein, crp1, is a smooth muscle marker. Developmental dynamics: an official publication of the American Association of Anatomists, 214(3):229–238, 1999.

[26] Thuan C Tran, CoreyAyne Singleton, Tamara S Fraley, and Jeffrey A Greenwood. Cysteine-rich protein 1 (crp1) regulates actin filament bundling. BMC cell biology, 6(1):45, 2005.

[27] David F Chang, Narasimhaswamy S Belaguli, Dinakar Iyer, Wilmer B Roberts, San-Pin Wu, Xiu-Rong Dong, Joseph G Marx, Mary Shannon Moore, Mary C Beckerle, Mark W Majesky, et al. Cysteine-rich lim-only proteins crp1 and crp2 are potent smooth muscle differentiation cofactors. Developmental cell, 4(1):107–118, 2003.

[28] Brenda Lilly, Kathleen A Clark, Masaaki Yoshigi, Stephen Pronovost, Meng-Ling Wu, Muthu Periasamy, Mei Chi, Richard J Paul, Shaw-Fang Yet, and Mary C Beckerle. Loss of the serum response factor cofactor, cysteine-rich protein 1, attenuates neointima formation in the mouse. Arteriosclerosis, thrombosis, and vascular biology, 30(4):694–701, 2010.

[29] Kota Y Miyasaka, Yasuyuki S Kida, Takayuki Sato, Mari Minami, and Toshihiko Ogura. Csrp1 regulates dynamic cell movements of the mesendoderm and cardiac mesoderm through interactions with dishevelled and diversin. Proceedings of the National Academy of Sciences, 104(27):11274–11279, 2007.

[30] Amina Kamar, Akl C. Fahed, Kamel Shibbani, Nehme El-Hachem, Salim Bou-Slaiman, Mariam Arabi, Mazen Kurban, Jonathan G. Seidman, Christine E. Seidman, Rachid Haidar, Elias Baydoun, Georges Nemer, and Fadi Bitar. A novel role for csrp1 in a lebanese family with congenital cardiac defects. Frontiers in Genetics, 8:217, 2017.

[31] Eran Eden, Roy Navon, Israel Steinfeld, Doron Lipson, and Zohar Yakhini. Gorilla: a tool for discovery and visualization of enriched go terms in ranked gene lists. BMC bioinformatics, 10(1):1–7, 2009.

[32] Molly A. Hall, Jason H. Moore, and Marylyn D. Ritchie. Embracing complex associations in common traits: Critical considerations for precision medicine. Trends in Genetics, 32(8):470–484, 2016.

[33] Michael D Gallagher and Alice S Chen-Plotkin. The post-gwas era: From association to function. Am J Hum Genet, 102(5):717–730, May 2018.

[34] Po-Ru Loh, Gaurav Bhatia, Alexander Gusev, Hilary K Finucane, Brendan K Bulik-Sullivan, Samuela J Pollack, Teresa R de Candia, Sang Hong Lee, Naomi R Wray, Kenneth S Kendler, Michael C O’Donovan, Benjamin M Neale, Nick Patterson, Alkes L Price, and Schizophrenia Working Group of the Psychiatric Genomics Consortium. Contrasting genetic architectures of schizophrenia and other complex diseases using fast variance-components analysis. Nature Genetics, 47(12):1385–1392, 2015.

[35] Kyle Kai-How Farh, Alexander Marson, Jiang Zhu, Markus Kleinewietfeld, William J. Housley, Samantha Beik, Noam Shoresh, Holly Whitton, Russell J. H. Ryan, Alexander A. Shishkin, Meital Hatan, Marlene J. Carrasco-Alfonso, Dita Mayer, C. John Luckey, Nikolaos A. Patsopoulos, Philip L. De Jager, Vijay K. Kuchroo, Charles B. Epstein, Mark J. Daly, David A. Hafler, and Bradley E. Bernstein. Genetic and epigenetic fine mapping of causal autoimmune disease variants. Nature, 518(7539):337–343, 2015.

[36] Matthew T. Maurano, Richard Humbert, Eric Rynes, Robert E. Thurman, Eric Haugen, Hao Wang, Alex P. Reynolds, Richard Sandstrom, Hongzhu Qu, Jennifer Brody, Anthony Shafer, Fidencio Neri, Kristen Lee, Tanya Kutyavin, Sandra Stehling-Sun, Audra K. Johnson, Theresa K. Canfield, Erika Giste, Morgan Diegel, Daniel Bates, R. Scott Hansen, Shane Neph, Peter J. Sabo, Shelly Heimfeld, Antony Raubitschek, Steven Ziegler, Chris Cotsapas, Nona Sotoodehnia, Ian Glass, Shamil R. Sunyaev, Rajinder Kaul, and John A. Stamatoyannopoulos. Systematic localization of common disease-associated variation in regulatory dna. Science, 337(6099):1190–1195, 2012.

[37] Jeff Vierstra, John Lazar, Richard Sandstrom, Jessica Halow, Kristen Lee, Daniel Bates, Morgan Diegel, Douglas Dunn, Fidencio Neri, Eric Haugen, Eric Rynes, Alex Reynolds, Jemma Nelson, Audra Johnson, Mark Frerker, Michael Buckley, Rajinder Kaul, Wouter Meuleman, and John A. Stamatoyannopoulos. Global reference mapping of human transcription factor footprints. Nature, 583(7818):729–736, 2020.

[38] Christian von Mering, Martijn Huynen, Daniel Jaeggi, Steffen Schmidt, Peer Bork, and Berend Snel. String: a database of predicted functional associations between proteins. Nucleic acids research, 31(1):258–261, 2003.

[39] Muredach P Reilly, Mingyao Li, Jing He, Jane F Ferguson, Ioannis M Stylianou, Nehal N Mehta, Mary Susan Burnett, Joseph M Devaney, Christopher W Knouff, John R Thompson, et al. Identification of adamts7 as a novel locus for coronary atherosclerosis and association of abo with myocardial infarction in the presence of coronary atherosclerosis: two genome-wide association studies. The Lancet, 377(9763):383–392, 2011.

[40] Martin Dichgans, Rainer Malik, Inke R König, Jonathan Rosand, Robert Clarke, Solveig Gretarsdottir, Gudmar Thorleifsson, Braxton D Mitchell, Themistocles L Assimes, Christopher Levi, et al. Shared genetic susceptibility to ischemic stroke and coronary artery disease: a genome-wide analysis of common variants. Stroke, 45(1):24–36, 2014.

[41] Majid Nikpay, Anuj Goel, Hong-Hee Won, Leanne M Hall, Christina Willenborg, Stavroula Kanoni, Danish Saleheen, Theodosios Kyriakou, Christopher P Nelson, Jemma C Hopewell, et al. A comprehensive 1000 genomes–based genome-wide association meta-analysis of coronary artery disease. Nature genetics, 47(10):1121, 2015.

[42] Salma M Wakil, Ramesh Ram, Nzioka P Muiya, Munish Mehta, Editha Andres, Nejat Mazhar, Batoul Baz, Samya Hagos, Maie Alshahid, Brian F Meyer, et al. A genome-wide association study reveals susceptibility loci for myocardial infarction/coronary artery disease in saudi arabs. Atherosclerosis, 245:62–70, 2016.

[43] Derek Klarin, Qiuyu Martin Zhu, Connor A Emdin, Mark Chaffin, Steven Horner, Brian J McMillan, Alison Leed, Michael E Weale, Chris CA Spencer, François Aguet, et al. Genetic analysis in uk biobank links insulin resistance and transendothelial migration pathways to coronary artery disease. Nature genetics, 49(9):1392, 2017.

[44] Pim van der Harst and Niek Verweij. Identification of 64 novel genetic loci provides an expanded view on the genetic architecture of coronary artery disease. Circulation research, 122(3):433–443, 2018.

[45] Yi Han, Rajkumar Dorajoo, Xuling Chang, Ling Wang, Chiea-Chuen Khor, Xueling Sim, Ching-Yu Cheng, Yuan Shi, Yih Chung Tham, Wanting Zhao, et al. Genome-wide association study identifies a missense variant at apoa5 for coronary artery disease in multi-ethnic cohorts from southeast asia. Scientific reports, 7(1):1–11, 2017.

[46] Yang Li, Dao Wen Wang, Yundai Chen, Can Chen, Jian Guo, Shu Zhang, Zhijun Sun, Hu Ding, Yan Yao, Lei Zhou, et al. Genome-wide association and functional studies identify scml4 and thsd7a as novel susceptibility genes for coronary artery disease. Arteriosclerosis, thrombosis, and vascular biology, 38(4):964–975, 2018.

[47] Wei Zhou, Jonas B Nielsen, Lars G Fritsche, Rounak Dey, Maiken E Gabrielsen, Brooke N Wolford, Jonathon LeFaive, Peter VandeHaar, Sarah A Gagliano, Aliya Gifford, et al. Efficiently controlling for case-control imbalance and sample relatedness in large-scale genetic association studies. Nature genetics, 50(9):1335–1341, 2018.

[48] Yoshiji Yamada, Yoshiki Yasukochi, Kimihiko Kato, Mitsutoshi Oguri, Hideki Horibe, Tetsuo Fujimaki, Ichiro Takeuchi, and Jun Sakuma. Identification of 26 novel loci that confer susceptibility to early-onset coronary artery disease in a japanese population. Biomedical reports, 9(5):383–404, 2018.

[49] John D Rioux, Ramnik J Xavier, Kent D Taylor, Mark S Silverberg, Philippe Goyette, Alan Huett, Todd Green, Petric Kuballa, M Michael Barmada, Lisa Wu Datta, et al. Genome-wide association study identifies new susceptibility loci for crohn disease and implicates autophagy in disease pathogenesis. Nature genetics, 39(5):596–604, 2007.

[50] Cécile Libioulle, Edouard Louis, Sarah Hansoul, Cynthia Sandor, Frédéric Farnir, Denis Franchimont, Séverine Vermeire, Olivier Dewit, Martine De Vos, Anna Dixon, et al. Novel crohn disease locus identified by genome-wide association maps to a gene desert on 5p13. 1 and modulates expression of ptger4. PLoS Genet, 3(4):e58, 2007.

[51] Miles Parkes, Jeffrey C Barrett, Natalie J Prescott, Mark Tremelling, Carl A Anderson, Sheila A Fisher, Roland G Roberts, Elaine R Nimmo, Fraser R Cummings, Dianne Soars, et al. Sequence variants in the autophagy gene irgm and multiple other replicating loci contribute to crohn’s disease susceptibility. Nature genetics, 39(7):830–832, 2007.

[52] Wellcome Trust Case Control Consortium et al. Genome-wide association study of 14,000 cases of seven common diseases and 3,000 shared controls. Nature, 447(7145):661, 2007.

[53] Andre Franke, Jochen Hampe, Philip Rosenstiel, Christian Becker, Florian Wagner, Robert Häsler, Randall D Little, Klaus Huse, Andreas Ruether, Tobias Balschun, et al. Systematic association mapping identifies nell1 as a novel ibd disease gene. PloS one, 2(8):e691, 2007.

[54] John V Raelson, Randall D Little, Andreas Ruether, Hélene Fournier, Bruno Paquin, Paul Van Eerdewegh, WEC Bradley, Pascal Croteau, Quynh Nguyen-Huu, Jonathan Segal, et al. Genome-wide association study for crohn’s disease in the quebec founder population identifies multiple validated disease loci. Proceedings of the National Academy of Sciences, 104(37):14747–14752, 2007.

[55] Jeffrey C Barrett, Sarah Hansoul, Dan L Nicolae, Judy H Cho, Richard H Duerr, John D Rioux, Steven R Brant, Mark S Silverberg, Kent D Taylor, M Michael Barmada, et al. Genome-wide association defines more than 30 distinct susceptibility loci for crohn’s disease. Nature genetics, 40(8):955–962, 2008.

[56] Dermot PB McGovern, Michelle R Jones, Kent D Taylor, Kristin Marciante, Xiaofei Yan, Marla Dubinsky, Andy Ippoliti, Eric Vasiliauskas, Dror Berel, Carrie Derkowski, et al. Fucosyltransferase 2 (fut2) non-secretor status is associated with crohn’s disease. Human molecular genetics, 19(17):3468–3476, 2010.

[57] Andre Franke, Dermot PB McGovern, Jeffrey C Barrett, Kai Wang, Graham L Radford-Smith, Tariq Ahmad, Charlie W Lees, Tobias Balschun, James Lee, Rebecca Roberts, et al. Genome-wide meta-analysis increases to 71 the number of confirmed crohn’s disease susceptibility loci. Nature genetics, 42(12):1118–1125, 2010.

[58] Jie Huang, David Ellinghaus, Andre Franke, Bryan Howie, and Yun Li. 1000 genomes-based imputation identifies novel and refined associations for the wellcome trust case control consortium phase 1 data. European Journal of Human Genetics, 20(7):801–805, 2012.

[59] Eimear E Kenny, Itsik Pe’er, Amir Karban, Laurie Ozelius, Adele A Mitchell, Sok Meng Ng, Monica Erazo, Harry Ostrer, Clara Abraham, Maria T Abreu, et al. A genome-wide scan of ashkenazi jewish crohn’s disease suggests novel susceptibility loci. PLoS Genet, 8(3):e1002559, 2012.

[60] Antonio Julià, Eugeni Domènech, Elena Ricart, Raül Tortosa, Valle García-Sánchez, Javier P Gisbert, Pilar Nos Mateu, Ana Gutiérrez, Fernando Gomollón, Juan Luís Mendoza, et al. A genome-wide association study on a southern european population identifies a new crohn’s disease susceptibility locus at rbx1-ep300. Gut, 62(10):1440–1445, 2013.

[61] Luke Jostins, Stephan Ripke, Rinse K Weersma, Richard H Duerr, Dermot P McGovern, Ken Y Hui, James C Lee, L Philip Schumm, Yashoda Sharma, Carl A Anderson, et al. Host–microbe interactions have shaped the genetic architecture of inflammatory bowel disease. Nature, 491(7422):119–124, 2012.

[62] Keiko Yamazaki, Junji Umeno, Atsushi Takahashi, Atsushi Hirano, Todd Andrew Johnson, Natsuhiko Kumasaka, Takashi Morizono, Naoya Hosono, Takaaki Kawaguchi, Masakazu Takazoe, et al. A genome-wide association study identifies 2 susceptibility loci for crohn’s disease in a japanese population. Gastroenterology, 144(4):781–788, 2013.

[63] Suk-Kyun Yang, Myunghee Hong, Wanting Zhao, Yusun Jung, Jiwon Baek, Naeimeh Tayebi, Kyung Mo Kim, Byong Duk Ye, Kyung-Jo Kim, Sang Hyoung Park, et al. Genome-wide association study of crohn’s disease in koreans revealed three new susceptibility loci and common attributes of genetic susceptibility across ethnic populations. Gut, 63(1):80–87, 2014.

[64] Suk-Kyun Yang, Myunghee Hong, Hyunchul Choi, Wanting Zhao, Yusun Jung, Talin Haritunians, Byong Duk Ye, Kyung-Jo Kim, Sang Hyoung Park, Inchul Lee, et al. Immunochip analysis identification of 6 additional susceptibility loci for crohn’s disease in koreans. Inflammatory bowel diseases, 21(1):1–7, 2015.

[65] Jimmy Z Liu, Suzanne Van Sommeren, Hailiang Huang, Siew C Ng, Rudi Alberts, Atsushi Takahashi, Stephan Ripke, James C Lee, Luke Jostins, Tejas Shah, et al. Association analyses identify 38 susceptibility loci for inflammatory bowel disease and highlight shared genetic risk across populations. Nature genetics, 47(9):979–986, 2015.

[66] Chengrui Huang, Talin Haritunians, David T Okou, David J Cutler, Michael E Zwick, Kent D Taylor, Lisa W Datta, Joseph C Maranville, Zhenqiu Liu, Shannon Ellis, et al. Characterization of genetic loci that affect susceptibility to inflammatory bowel diseases in african americans. Gastroenterology, 149(6):1575–1586, 2015.

[67] Eun Suk Jung, Jae Hee Cheon, Ji Hyun Lee, Soo Jung Park, Hui Won Jang, Sook Hee Chung, Myoung Hee Park, Tai-Gyu Kim, Heung-Bum Oh, Suk-Kyun Yang, et al. Hla-c* 01 is a risk factor for crohn’s disease. Inflammatory bowel diseases, 22(4):796–806, 2016.

[68] Jerzy Ostrowski, Agnieszka Paziewska, Izabella Lazowska, Filip Ambrozkiewicz, Krzysztof Goryca, Maria Kulecka, Tomasz Rawa, Jakub Karczmarski, Michalina Dabrowska, Natalia Zeber-Lubecka, et al. Genetic architecture differences between pediatric and adult-onset inflammatory bowel diseases in the polish population. Scientific reports, 6:39831, 2016.

[69] Katrina M De Lange, Loukas Moutsianas, James C Lee, Christopher A Lamb, Yang Luo, Nicholas A Kennedy, Luke Jostins, Daniel L Rice, Javier Gutierrez-Achury, Sun-Gou Ji, et al. Genome-wide association study implicates immune activation of multiple integrin genes in inflammatory bowel disease. Nature genetics, 49(2):256–261, 2017.

[70] Yoichi Kakuta, Yosuke Kawai, Takeo Naito, Atsushi Hirano, Junji Umeno, Yuta Fuyuno, Zhenqiu Liu, Dalin Li, Takeru Nakano, Yasuhiro Izumiyama, et al. A genome-wide association study identifying rap1a as a novel susceptibility gene for crohn’s disease in japanese individuals. Journal of Crohn’s and Colitis, 13(5):648–658, 2019.

