## Supplementary Material for "Predicting genotype-specific gene regulatory networks"

### Supplementary Material:
#### Predicting genotype-specific gene regulatory networks

**Deborah Weighill, Marouen Ben Guebila, Kimberly Glass, John Quackenbush, John Platig**

#### Note S1 Reference motif prior

The hg19 human reference assembly was scanned for the presence of TF motifs using FIMO [1] and applying a p-value cutoff of $10^{-4}$. Motifs that were present within the promoter regions of genes were selected by identifying motifs that overlapped with the 1kb region (-750, +250) around all possible transcription start sites (TSS) of a gene (i.e. the TSS of each transcript of a gene), making use of the GenomicRanges R package [2]. Transcription start sites for each transcript were downloaded from the UCSC Table Browser https://genome.ucsc.edu/cgi-bin/hgTables, in the Ensembl genes table for hg19 on 06/10/2020. The resulting mapped motifs were then collapsed to construct the reference motif prior network $M$ defined as:

$$M_{ij} = \begin{cases} 1 & \text{if motif of TF } i \text{ overlaps with promoter region of gene } j \\ 0 & \text{otherwise} \end{cases}$$

#### Note S2 eQTLs, genotypes and QBiC

Expression QTLs for LCLs from GTEx version 7 [3] were downloaded from https://gtexportal.org/home/datasets on 06/10/2020. These eQTLs were then filtered to select only eQTLs where the variant resided within a TF motif within a promoter region (described in Note S2) *and* where the eGene was the gene adjacent to (and associated with) the promoter. Genotypes for NA12878 (corresponding to the GM12878 cell line) and K562 were downloaded on 06/10/2020. The Platinum Genomes genotype for NA12878 was obtained from https://www.illumina.com/platinumgenomes.html and the K562 genotype was obtained from ENCODE https://www.encodeproject.org/files/ENCFF538YDL/ derived from a study by Zhou *et al.* (2019) [4]. Using the eQTL variants within motifs, we selected those variants where at least one of the cell lines (K562 or GM12878) had at least one alternate allele of the eQTL variant. QBiC [5] was then run on these eQTLs, using hg19 as a reference genome.

#### Note S3 Prior modification

When running EGRET, a genotype-specific prior ("EGRET prior") is constructed for each individual. For each SNP within a given individual, the alternate allele count of the individual is calculated. For each eQTL variant $s$ in promoter region of gene $j$ within a motif for TF $i$, three attributes are assigned: (1) the alternate allele count of the individual at that location $A_{s_{ij}}$ (2) the beta value of the eQTL $\beta_{s_{ij}}$ and (3) the QBiC effect of the SNP $q_{s_{ij}}$ on the binding of the TF corresponding to the motif in which the variant resides (only significant negative QBiC values are used). The effect of a SNP on TF binding in the given individual is then defined as the product $|q_{s_{ij}} A_{s_{ij}} \beta_{s_{ij}}|$. Modifier weights to the reference motif prior are then calculated by aggregating these effects per TF-gene pair, allowing for the fact that a gene might have more than one variant in its promoter region affecting the binding of a particular TF. The genotype-specific prior edge weight $E_{ij}$ for TF $i$ and gene $j$ is thus defined as

$$E_{ij} = M_{ij} - \sum_{s} |q_{s_{ij}} A_{s_{ij}} \beta_{s_{ij}}|$$

where $M_{ij}$ is the reference motif prior defined above in Note S1.

#### Note S4 Gene expression and PPI data

Gene expression data as TPMs (transcripts per million) for lymphoblastoid cell lines (LCLs) from The Genotype-Tisse Expression Project (GTEx) version 7 [3], was downloaded from https://gtexportal.org/home/datasets on 06/10/2020. The expression matrix was pruned to keep only genes that had non-zero expression values in at least 50 samples. The protein-protein interaction network as used in [6] was then filtered to keep only proteins whose corresponding genes met the same expression requirements described above (non-zero expression values in at least 50 samples). Thus, when selecting the set of genes and TFs to be included in the GRN, we removed any TFs or genes that did not have reasonable evidence of expression, where we defined "reasonable evidence of expression" as having non-zero values in ≥50 samples. Gene ID mapping from TF gene names to ensembl IDs was done using the mapping downloaded from ftp://ftp.ensembl.org/pub/grch37/current/gtf/homo_sapiens/Homo_sapiens.GRCh37.87.chr.gtf.gz on 06/10/2020.

#### Note S5 Message Passing

##### Note S5.1 The PANDA Message Passing Framework

"Passing Attributes between Networks for Data Assimilation" (PANDA) is a framework for gene regulatory network (GRN) construction [7], that uses a message passing approach to combine (1) transcription factor (TF) motif information, (2) gene co-expression, and (3) TF protein-protein interactions (PPI) to estimate a bipartite GRN modeling the regulatory interactions between TFs and genes. The message passing process finds agreement between these three data types, each represented as a network, and produces a final GRN based on evidence of gene regulation from each of these data layers.

PANDA begins with a reference motif network $M$ representing a preliminary estimate of potential regulatory relationships between TFs and genes based on TF motif information, and is defined above in Note S1.

PANDA also uses two other sources of regulatory information, namely a gene co-expression network $C$ representing potential co-regulatory relationships between genes, and a PPI network $P$ representing which TFs may physically interact to form protein complexes. PANDA then uses message passing to iteratively calculate the similarity between these networks and incrementally update the network edge weights to reflect information from the other networks. For each TF-gene pair $ij$ in $M$, the *availability* of the edge represents the similarity between the target genes of TF $i$ in $M$ and the set of genes with which gene $j$ is co-expressed in $C$. For the same edge in $M$, the *responsibility* represents the similarity between the set of TFs that target gene $j$ in $M$ and the interaction partners of TF $i$ in $P$. Edge weights in $M$ are then updated with a small fraction ($\alpha = 0.1$) of the average of the *responsibility* and *availability*. Each edge in $C$ and $P$ is also updated with a small fraction of the overlap of the neighbors of pairs of genes/TFs in $M$, respectively. For example, the edge $kl$ in $P$ will be updated with the similarity between the set of target genes of TFs $k$ and $l$ in $M$. These updates of $M$, and then $C$ and $P$ are iteratively repeated until convergence is achieved.

PANDA has been applied in the investigation of GRN patterns in several diseases including chronic obstructive pulmonary disease [8], asthma [9], ovarian cancer [10], and colon cancer [11]. In addition, PANDA has been used to study how gene regulation varies across different tissues [6, 12].

##### Note S5.2 Message Passing in EGRET

We used the PANDA message passing framework [7] in the pandaR package to combine the the EGRET prior $E$, PPI prior $P$, and gene expression data $C$, resulting in a predicted genotype-specific gene regulatory network for an individual. This process was applied to both GM12878 and K562 genotypes. Parameters used in the message passing were: remove.missing.ppi = TRUE, remove.missing.motif = TRUE, remove.missing.genes = TRUE. These parameters ensure that the set of TFs is defined by those in the motif-gene prior, and that the set of genes is defined as the intersection of those in the motif-gene prior and the gene expression matrix. In addition, a "genotype agnostic" baseline GRN ($B^*$) was constructed using the PANDA framework, running message passing on $M$, $P$, and $C$. This provided a baseline GRN for comparison with the genotype-specific GRNs.

#### Note S6 Comparison of EGRET networks from two cell line genotypes

##### Note S6.1 ChIP-seq regulatory network

ChIP-seq data from ReMap2018 for GM12878 and K562 [13] (hg19 reference genome) was downloaded from `http://pedagogix-tagc.univ-mrs.fr/remap/index.php?page=download` on 06/12/2020. This consisted of genomic ranges in BED format corresponding to the identified binding positions of several transcription factors (110 TFs for GM12878 and 204 TFs for K562). From this ChIP-seq data, TF binding sites within the promoter regions of genes were selected in the same manner as as the motif regions, described above. This resulted in two validation networks $V$, one for each cell line, where

$$V_{ij} = \begin{cases} 1 & \text{if ChIP-seq range of TF } i \text{ overlaps with promoter region of gene } j \\ 0 & \text{otherwise} \end{cases}$$

The subset of TFs for which ChIP-seq data was available in a given genotype (GM12878 or K562) were then used for subsequent analysis involving comparison of EGRET networks with ChIP-seq networks.

##### Note S6.2 Improving prediction of TF binding

The top edges with the highest disruption scores $d_{x_{ij}}^{(E)}$ were selected from the EGRET GM12878 and K562 networks, using a selection of different $d_{x_{ij}}^{(E)}$ cutoffs to define the top set of edges (Tables S3 and S4). Using the EGRET edge score as the predictor variable and the edges from the gold standard ChIP-seq GRN $V$ as the ground truth, we calculated performance metrics, namely the area under the receiver-operator characteristic (AU-ROC) and the area under the precision-recall (AU-PR) curve for edges with the top disruption scores $d_{x_{ij}}^{(E)}$; this was repeated for different thresholds of the edge disruption score. To compare the EGRET edge weights with those from the genotype-agnostic network, we calculated the significance between the differences of the AUCs using the Delong test for comparing AUCs (Tables S3 and S4). In both GM12878 and K562, the genotype-specific edges significantly improved the prediction of TF binding on variant-impacted edges. An optimal threshold of $d_{x_{ij}}^{(E)} \geq 0.35$ was identified for the isolation of variant impacted edges, as this was the threshold at which ChIP-seq TF binding predictions improved significantly for both GM12878 and K562. AU-ROCs and AU-PRs were calculated using the precrec [14] and pROC [15] R packages.

##### Note S6.3 Allele-specific expression

Allele-specific expression (ASE) data using the BiT-STARR-seq method in LCLs [16] was downloaded from `https://genome.cshlp.org/content/suppl/2018/10/17/gr.237354.118.DC1/Supplemental_Table_S1_.txt` on 09/01/2020. This data contained all variants tested for an ASE association, and these variants were mapped to TF motif regions in the promoter regions of genes, in the same manner as described above. Each gene $j$ was then assigned *gene regulatory difference score* $R_j^{(G)}$ defined as:

$$R_j^{(G)} = \sum_i R_{ij}^{(E)}.$$

This score agglomerates the *edge regulatory difference scores* per gene, providing a metric quantifying the total extent to which a gene's promoter region is differentially disrupted between the two cell lines. We then used Fisher's exact test to determine whether genes with a high regulatory difference score $R_j^{(G)}$ between the two genotypes K562 and GM12878 were enriched for genes having a significant (FDR $\leq$ 0.1) ASE variant within a motif in their promoter region. "High" regulatory difference scores were considered to be those in to top 10%.

##### Note S6.4 Chromatin accessibility QTLs

Chromatin accessibility QTLs (caQTLs) determined in lymphoblastoid cell lines (LCLs) [17] were downloaded from http://eqtl.uchicago.edu/yri_ipsc/cht_results_full_LCL.txt on 09/08/2020 [17]. This data set contained all variants tested for a caQTL association. We mapped these variants to TF motif regions in the promoters of genes (described above in Note S2). We again used the regulatory difference scores $R_j^{(G)}$ to test whether genes with a high regulatory difference score $R_j^{(G)}$ between the two genotypes K562 and GM12878 were enriched for genes having a significant (FDR $\leq$ 0.1) caQTL variant within a motif in their promoter region, using Fisher's exact test in a similar manner as described above.

#### Note S7 Population study of 119 individuals across 3 cell types

##### Note S7.1 Network Construction

Gene expression and eQTL data for a population of lymphoblastoid cell lines (LCL), induced pluripotent stem cells (iPSCs) and cardiomyocytes (CMs) that were differentiated from the induced pluripotent stem cells derived the study by Banovich *et al.* [17] and Li *et al.* [18], as well as the corresponding genotypes of 119 Yoruba individuals were downloaded on 06/17/2020. Expression data and eQTLs for LCLs, as well as eQTLs for iPSCs and iPSC-CMs were downloaded from http://eqtl.uchicago.edu/ whereas gene expression data for iPSCs and CMs were obtained through the Gene Expression Omnibus (GEO) from https://www.ncbi.nlm.nih.gov/geo/query/acc.cgi?acc=GSE107654. For each cell type, significant eQTLs (p $\leq$ 1e-5) for genes where the SNP resided within a TF motif within the promoter region of a gene ([-750,+250] around a TSS) were selected.

For each cell type, SNPs in the population of 119 Yoruba individuals that also were selected as eQTLs in the respective cell type were then isolated, and QBiC was run on this set of SNPs, per cell type, as in Note S3.

LCL and iPSC expression data were already preprocessed through WASP and normalized by standardizing by gene and quantile normalizing by individual, a method developed and used in [19]. The CM expression was not yet normalized, and we followed the process detailed in the series matrix files from GEO https://www.ncbi.nlm.nih.gov/geo/query/acc.cgi?acc=GSE107654 in order to process the CM expression data in the same manner. This involved scaling each gene by mean centering and dividing by the standard deviation, followed by quantile-normalizing the individuals using the normalize.quantiles function in the preprocessCore R package [20]. QBiC [5] was then run on the eQTLs to predict the effect these SNPs had on the binding of TFs using the full set of TF binding models in QBiC, and using hg19 as a reference genome.

EGRET was then run for each genotype in each cell type (a total of $119 \times 3 = 357$ EGRET runs). In addition, message passing was performed using the co-expression network, PPI network, and the reference motif prior (which involves no genotype information) to construct a "genotype agnostic" baseline GRN for each cell type. Message passing was performed using the pandaR R package [7] and run in parallel using GNU Parallel [21].

##### Note S7.2 TF disruption scores

Edge disruption scores $d_{x_{ij}}^{(E)}$ described above were calculated for each edge in each individual network for each cell type, and thresholded at a value of 0.35. Subsequently, a TF disruption score $d_{x_i}^{(TF)}$ was calculated for each TF as

$$d_{x_i}^{(TF)} = \sum_j \left| E^*_{x_{ij}} - B^*_{ij} \right|$$

where $E^*_{x_{ij}}$ is the genotype-specific EGRET edge weight (after message passing) between TF $i$ and gene $j$, and $B^*_{ij}$ is the baseline GRN edge weight between TF $i$ and gene $j$. A scaled TF disruption score $d_{x_i}^{(TF)'}$ for a TF within in an individual and cell type was then calculated by subtracting the mean TF disruption score for that individual/cell type and dividing by the standard deviation. Disease-associated genes for coronary artery disease (CAD) and Crohn's disease (CD) were obtained from the GWAS catalog at `https://www.ebi.ac.uk/gwas/api/search/downloads/full` on 06/30/2020 [22]. See Tables S5 and S6 for a complete list of citations for the individual GWAS studies from which summary statistics were used.

##### Note S7.3 Differential modularity with ALPACA

For each individual, we used ALPACA [23] to compare the modularity of the individual's genotype-specific EGRET GRN with the baseline GRN, resulting in a score for each node representing the contribution of that node to the differential modularity. These scores were then quantile normalized per individual per cell type. Following that, scores were normalized by gene, first by mean-centering and then by scaling to standard deviation of one.

##### Note S7.4 GO Enrichment

Gene ontology enrichment was performed using GORILLA [24] on the web server available at `http://cbl-gorilla.cs.technion.ac.il/` using the "Single ranked list of genes " option and a p-value threshold of $10^{-3}$.

#### Note S8 Importance of data types

We ran a selection of EGRET versions in order to investigate the contributions of different data types that EGRET uses. First, we ran EGRET using data for GM12878 leaving out the gene expression and PPI information in the message passing, and assessed the ability of the resulting EGRET edge weights to predict GM12878-derived ChIP-seq-derived TF binding for the subset of the network where ChIP-seq data was available, using the ChIP-seq regulatory network described in Note S6.1. EGRET edge weights using all data types showed almost 5% improvement in ROC-AUC for predicting the gold standard ChIP-seq network when compared to the EGRET network run without expression and PPI data in the message passing.

We also tested versions of EGRET that left out data types from the prior modification. For example, we ran a version of EGRET leaving out QBiC effects, thus only requiring genetic variants to be eQTLs and not requiring those variants to have significant QBiC effects. Similarly, we ran a version leaving out eQTL data, only requiring variants in an individual to have a significant QBiC effect, and not requiring the variants to be eQTLs. In validating the ability of disrupted edges ($d_{x_{ij}}^{(E)} \geq 0.35$) to predict the corresponding ChIP-seq edges, we found that all data types were crucial in improving the accuracy of prediction, and that leaving out either eQTLs or QBiC effects was detrimental to the validity of the network structure (Figure S11).

#### Supplementary Figures

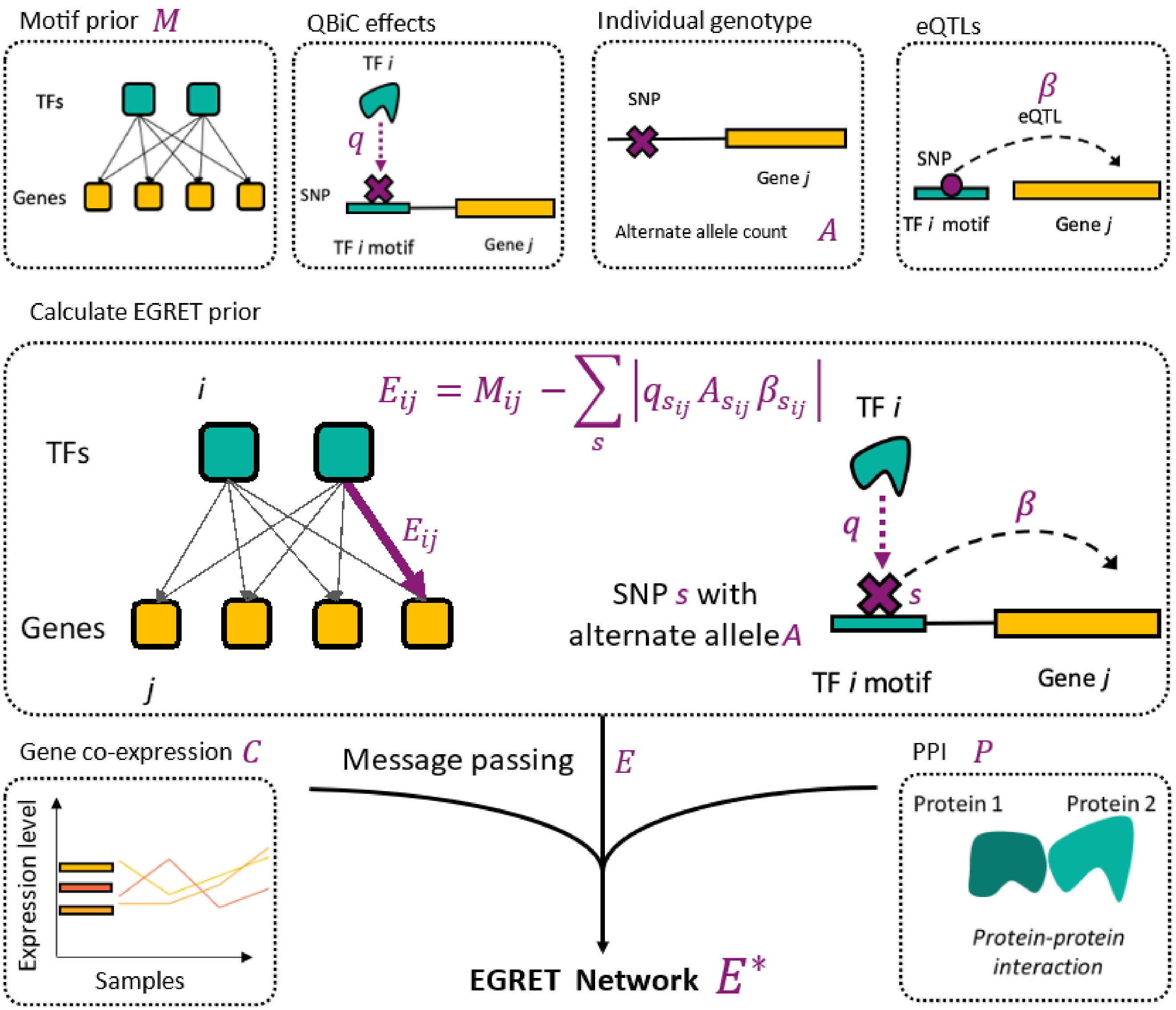


Figure S1: **Diagram illustrating the process and datatypes required for EGRET network construction.** EGRET begins with a reference motif prior representing the presence/absence of TF motifs in the promoter regions of genes. This is then modified by the individual's genetic mutations, penalizing motif-gene edges in which there exists a variant within the TF motif for which the individual has the alternate allele ($A$), the variant is an eQTL for the adjacent gene ($\beta$) and the variant is predicted through QBiC to disrupt TF binding at that location ($q$). These prior edges are then penalized by the absolute value of the product of the alternate allele count, the QBiC effect, and the eQTL beta value. Message passing then integrates the co-expression network ($C$) and PPI network ($P$) with the EGRET prior ($E$), resulting in a final genotype-specific GRN per individual ($E^*$).

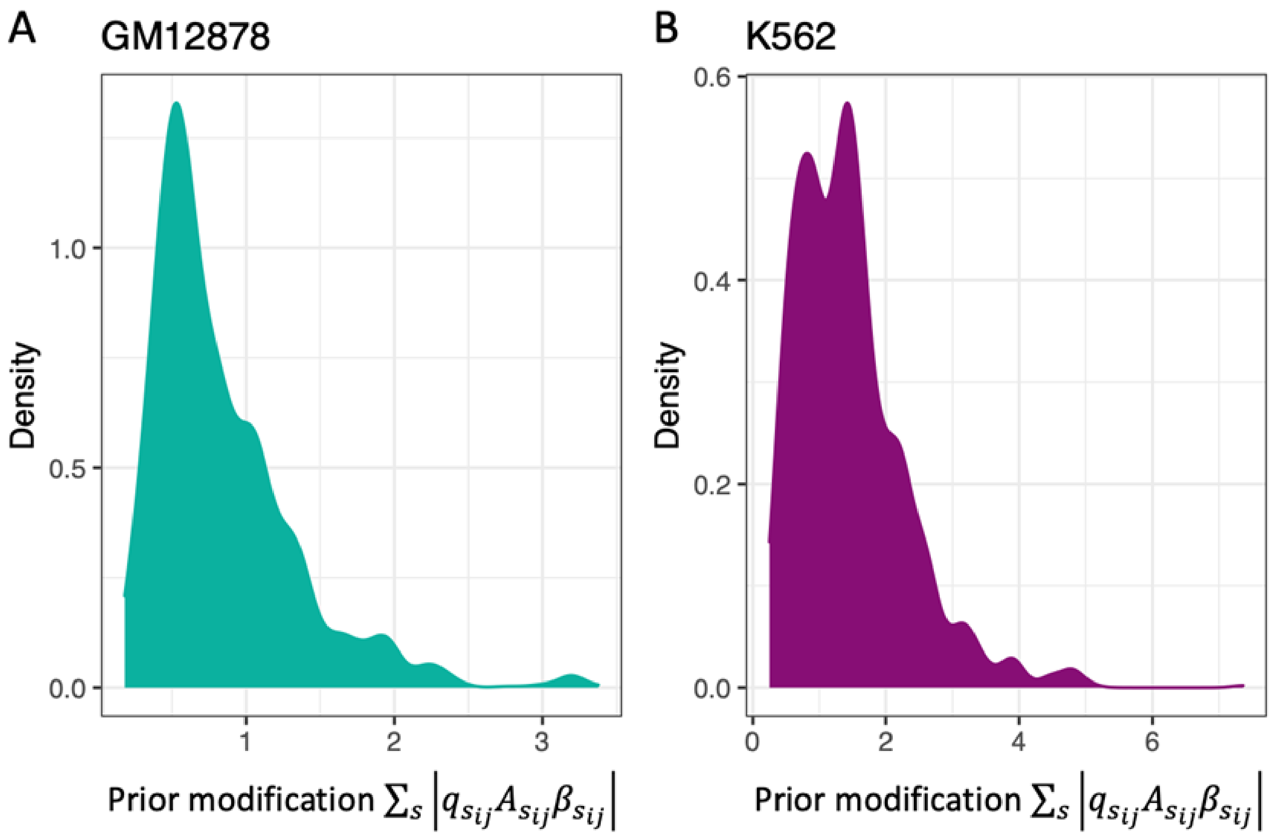


Figure S2: **Distribution of non-zero prior modifications** $\sum_s |q_{s_{ij}} A_{s_{ij}} \beta_{s_{ij}}|$ for (**A**) GM12878 and (**B**) K562.

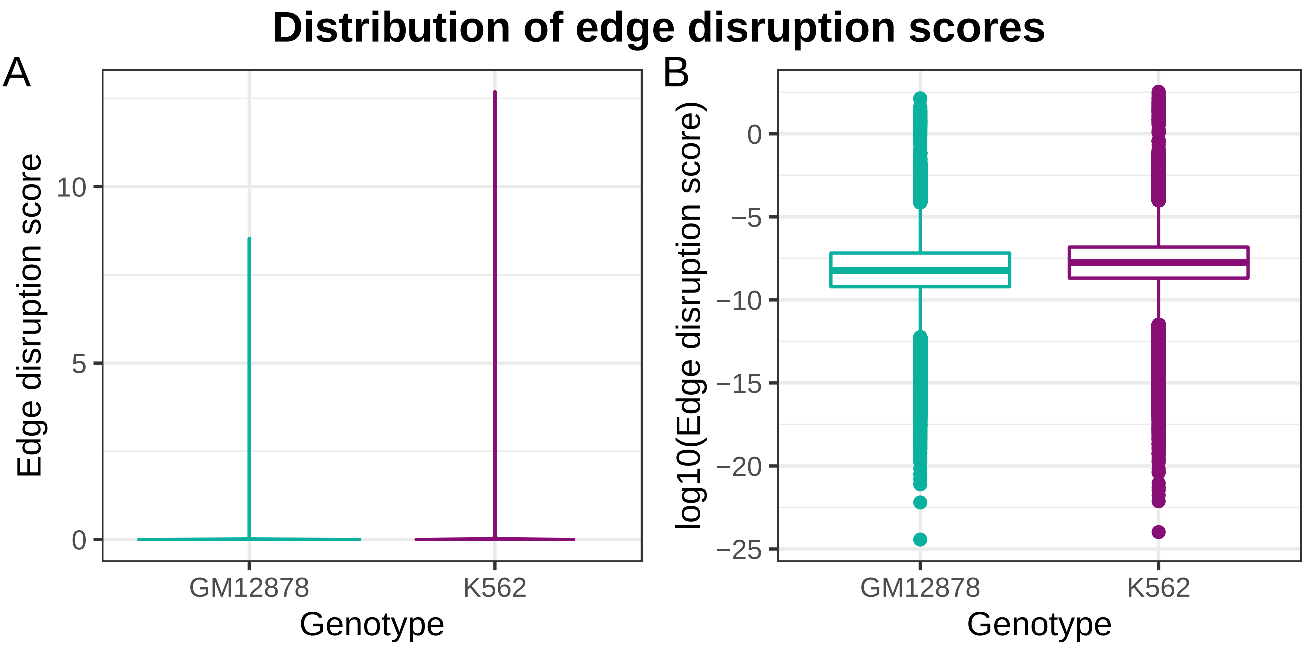


Figure S3: **Distribution of edge disruption scores** $d^{(E)}_{x_{ij}}$ **for GM12878 and K562.** (**A**) Violin plot of edge disruption scores. (**B**) Boxplot of $\log_{10}$ disruption scores.

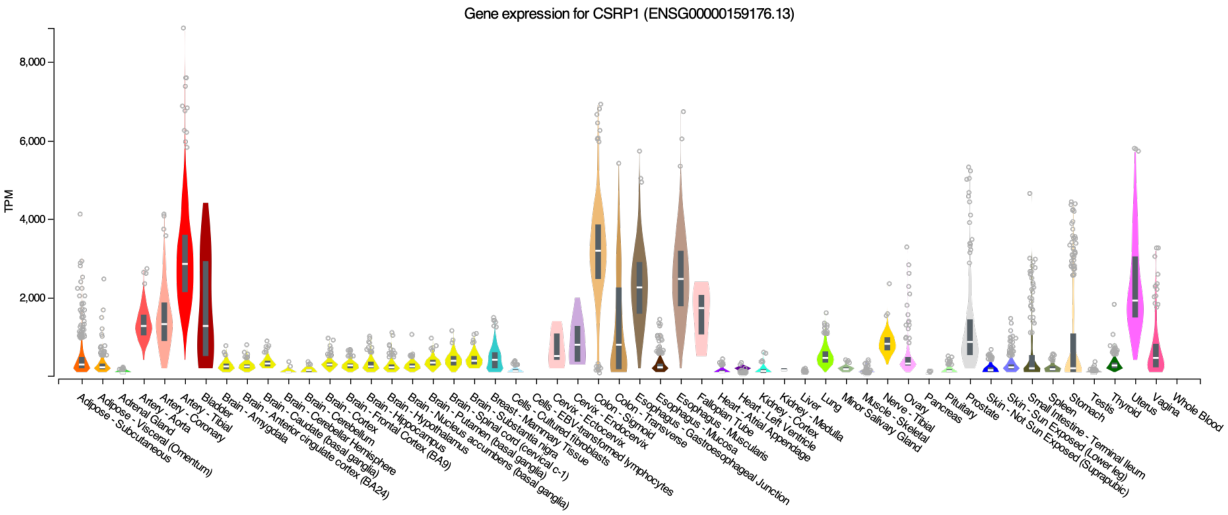


Figure S4: **CSRP1 expression.** TPM expression level of CSRP1 (ENSG00000159176) across all tissues available in GTEx. Plot obtained from the GTEx portal [3].

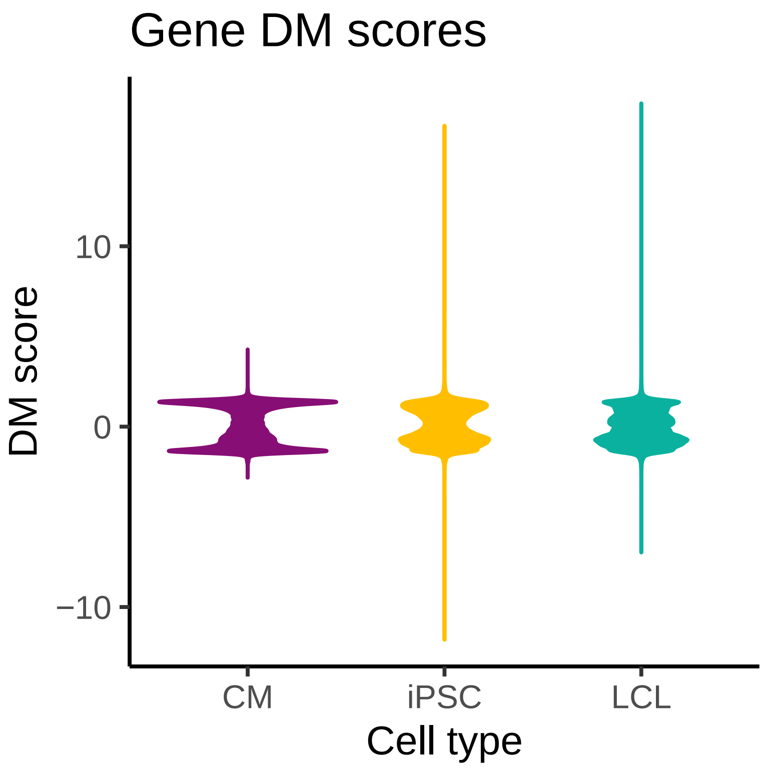


Figure S5: **Differential modularity scores.** Distributions of scaled differential modularity (DM) scores of genes in EGRET networks from 119 Yoruba individuals in three cell types.

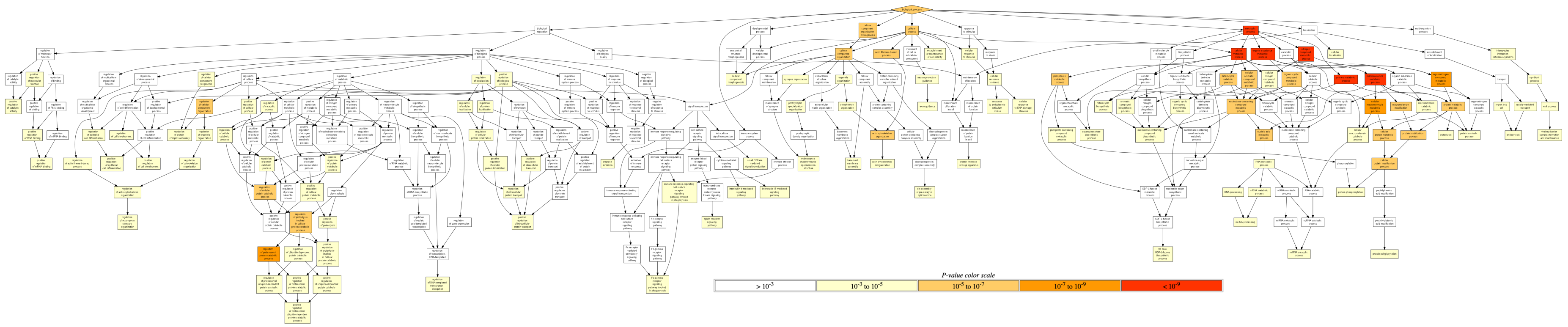


Figure S6: **Hierarchy of GO terms enriched in the genes with highest DM scores in individual 18 (genotype NA18523) in CMs, based on the mHG score test [24]. Enrichment performed using GORILLA [24].** GO terms are colored according to the significance of the p-value.

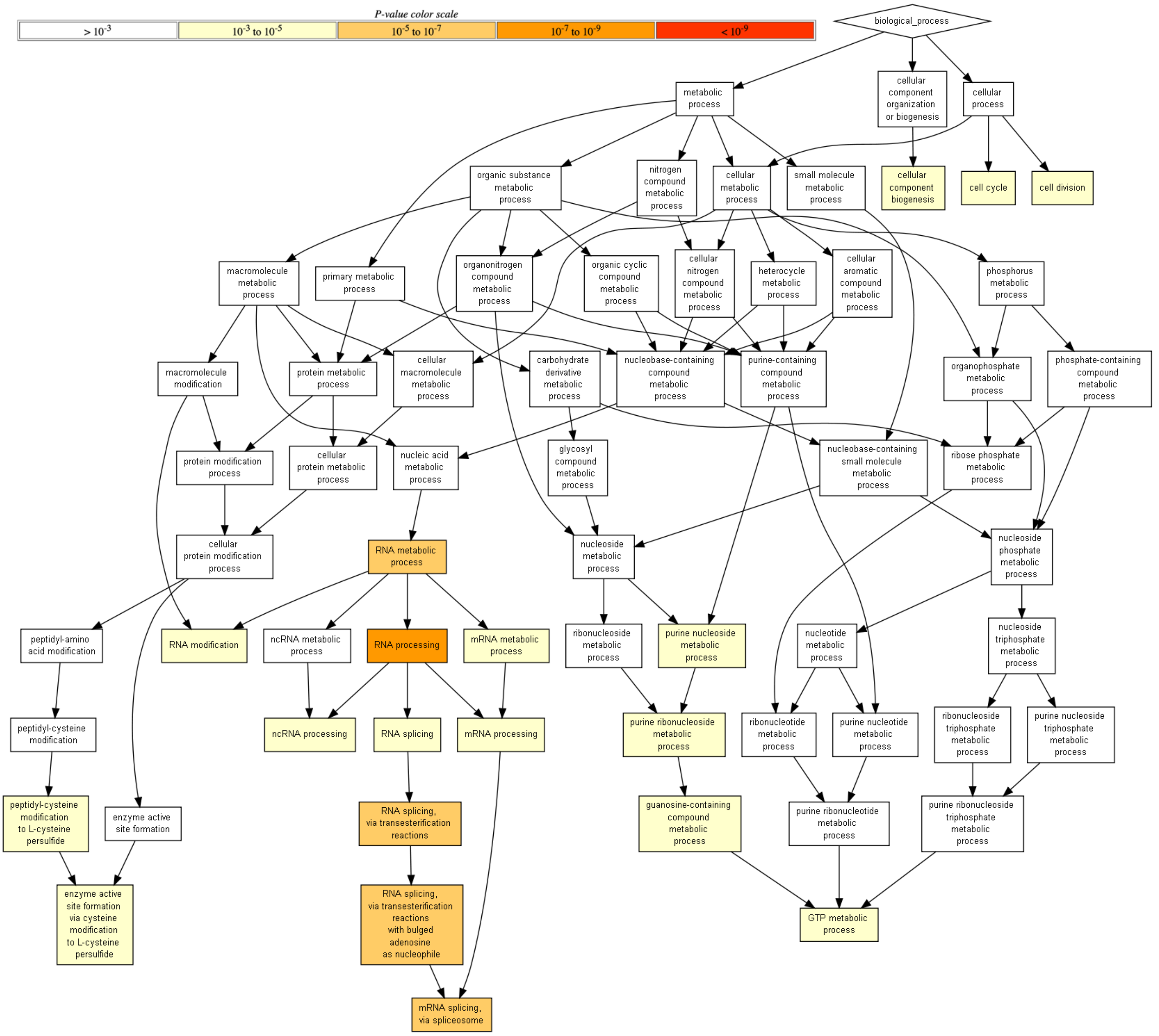


Figure S7: **Hierarchy of GO terms enriched in the genes with highest DM scores in individual 18 (genotype NA18523) in iPSCs, based on the mHG score test [24].** Enrichment performed using GORILLA [24]. GO terms are colored according to the significance of the p-value.

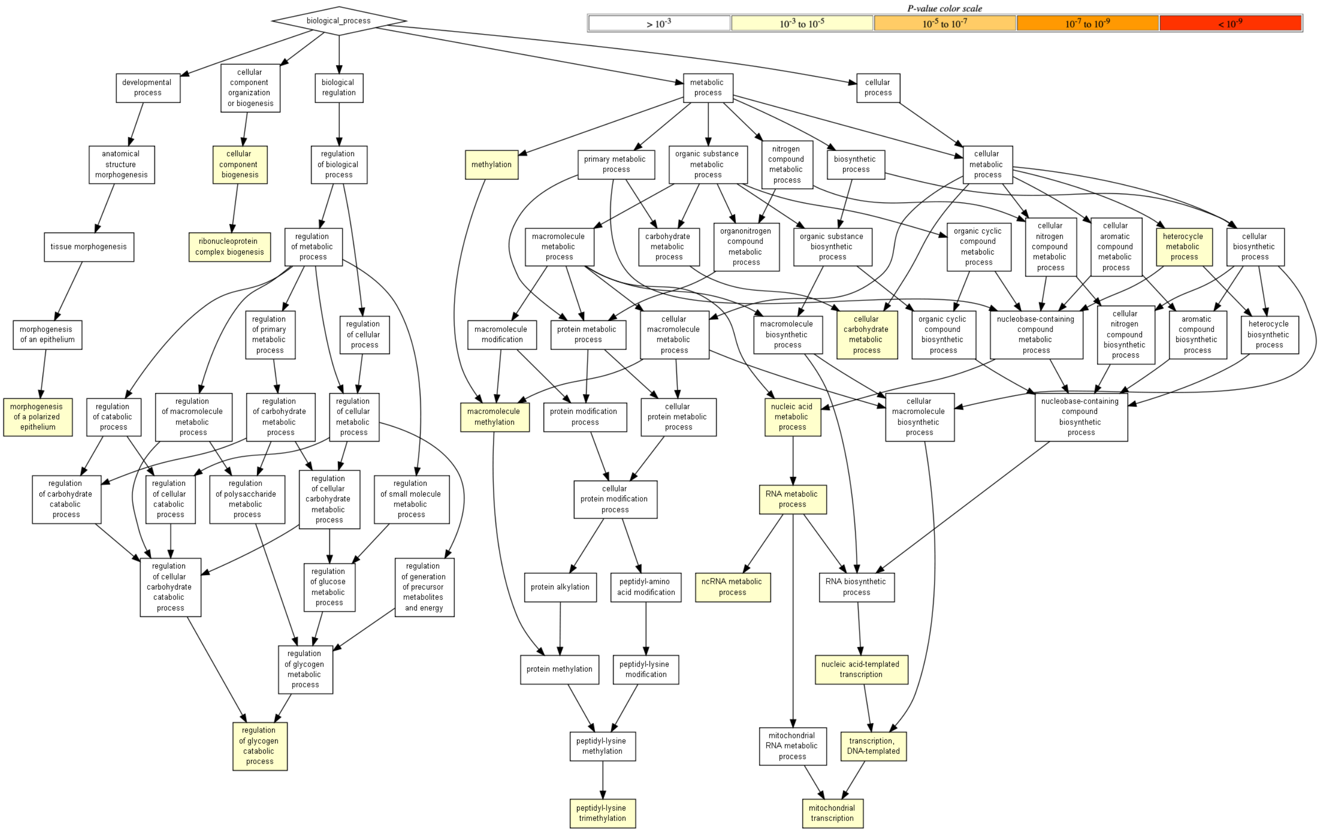


Figure S8: **Hierarchy of GO terms enriched in the genes with highest DM scores in individual 18 (genotype NA18523) in LCLs, based on the mHG score test [24].** Enrichment performed using GORILLA [24]. GO terms are colored according to the significance of the p-value.

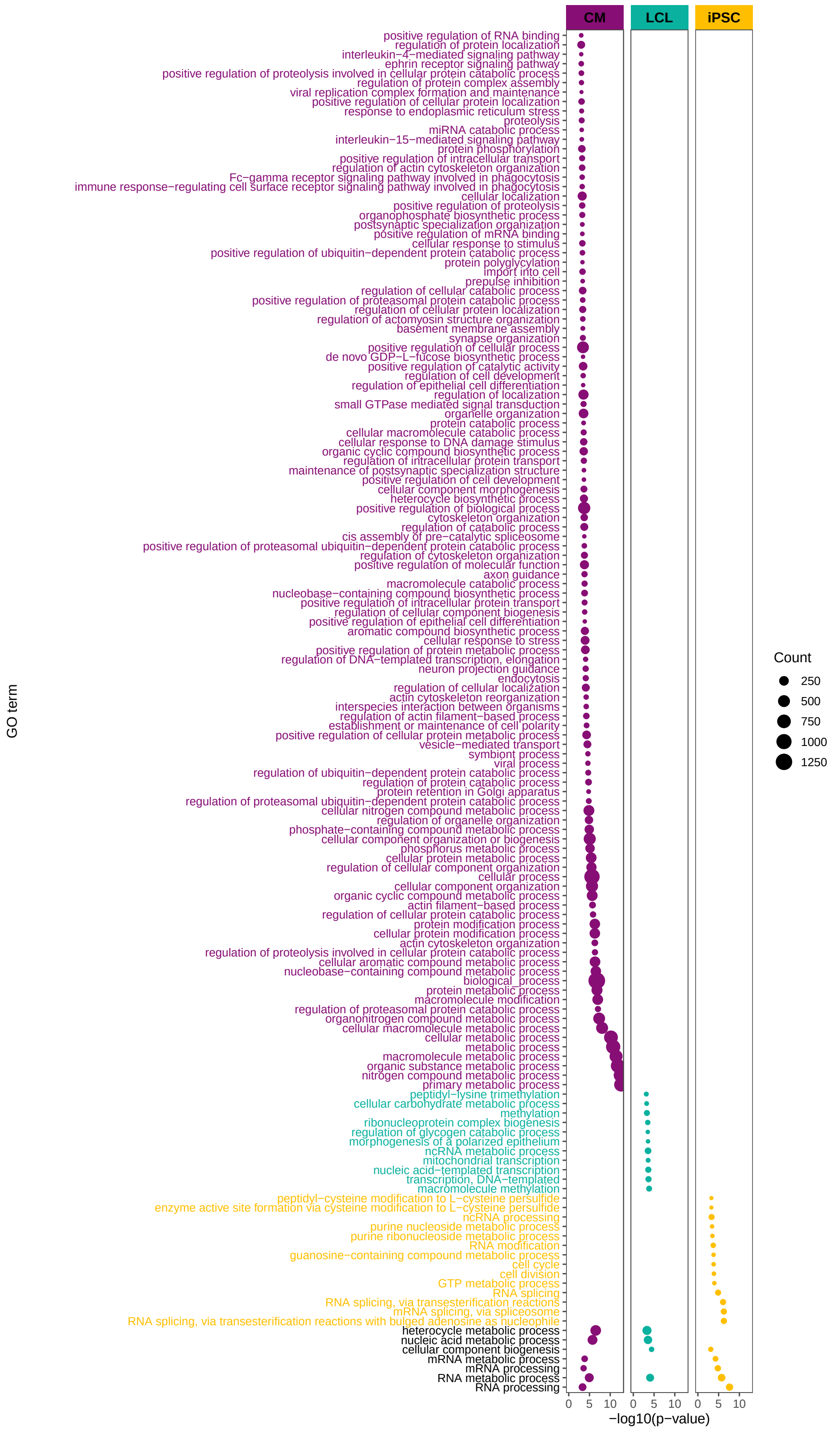


Figure S9: **GO terms enriched in genes with high DM scores of individual 18**, the individual with the highest TF disruption score for ERG. Point size corresponds to the the number of high-DM genes annotated with the corresponding GO term.

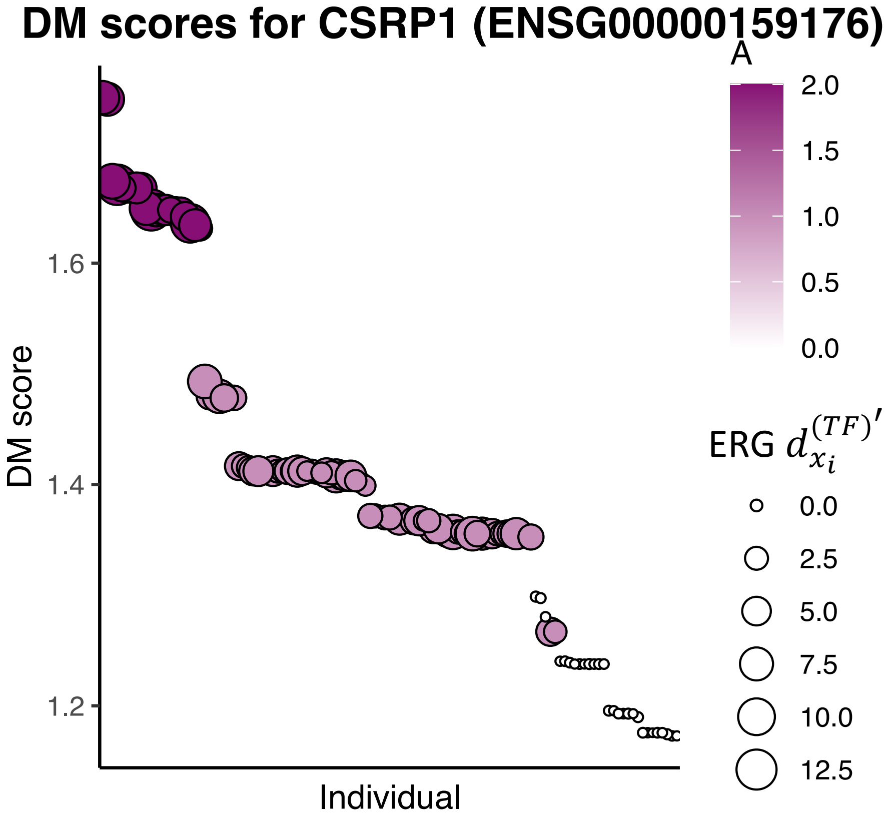


Figure S10: **DM scores of CSRP1 for 119 individuals in Yoruba population.** Point color intensity corresponds to the individual's alternate allele dosage $A$ for the SNP within the ERG binding motif, and point size corresponds to the TF disruption $d_{x_i}^{(TF)'}$ score of ERG.

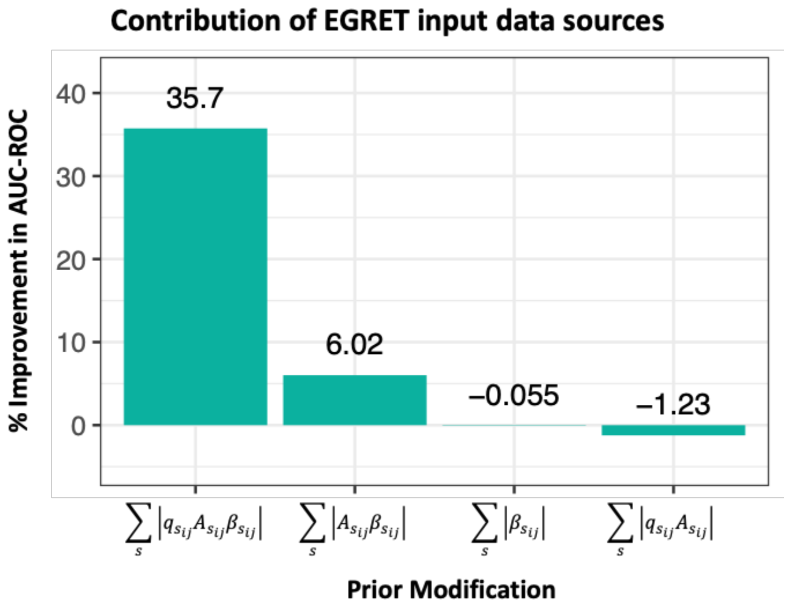


Figure S11: **Contribution of different data types to EGRET.** Percentage improvement in the prediction of the ChIP-seq regulatory network by the EGRET network $E^*$ in GM12878, compared to that of the baseline network $B^*$. Each bar represents the AUC-ROC improvement when using a different combination of data types in the prior modification, for each SNP $s$ with QBiC effect $q$, alternate allele count $A$ and eQTL beta value $\beta$. Percentage improvement calculated as $(AUC_{E^*} - AUC_{B^*})/AUC_{B^*}$

#### Supplementary Tables

Table S1: Inputs to EGRET.

| Name | Symbol | Source | Notes |
|---|---|---|---|
| Reference motif prior | $M$ | FIMO [1] motif calls | TF motif locations called on a reference genome. |
| eQTLs | $\beta$ | GTEx [3] | eQTLs from a public database can be used, it is not necessary for eQTLs to be determined in the investigator's specific study population. |
| SNPs | $s$ | Individual(s) genotype(s) | Genotypes of the specific individuals for which the investigator wishes to construct EGRET networks for. These are the variants used to tailor the networks to a specific individual. |
| Individual(s) | $x$ | Investigator's study individual/population | This is the individual(s) for which genotype-specific EGRET networks will be constructed. |
| QBiC predictions | $q$ | QBiC [5] | QBiC must be applied to the SNPs $s$ from the individual(s) $x$ which overlap with eQTLs within promoter-residing motifs. |
| PPI | $P$ | StringDB [25] | Protein-protein interactions from a public database. |
| Gene expression | $C$ | GTEx [3], TCGA | RNA-seq measurements across a population are needed to estimate co-expression relationships. $C$ does not need to be estimated from individuals for which specific networks are being constructed, it can be derived from databases such as GTEx and TCGA. |

Table S2: Calculated outputs from EGRET.

| Name | Symbol | Source/Formula | Notes |
|---|---|---|---|
| Egret prior | $E$ | $M - \sum_s \lvert q_{s_{ij}} A_{s_{ij}} \beta_{s_{ij}} \rvert$ | EGRET prior $E$ is the genotype-edited form of $M$ and is combined with $C$ and $P$ during message passing. |
| EGRET GRN | $E^*$ | Message passing | Genotype-specific EGRET GRN $E^*$ produced by message passing of $E$, $C$, and $P$. A high edge weight indicates a putative regulatory relationship where as a low weight indicates lack of a regulatory relationship. |
| Baseline GRN | $B^*$ | Message passing | Baseline GRN produced by message passing of $M$, $C$, and $P$. A high edge weight indicates a putative regulatory relationship where as a low weight indicates lack of a regulatory relationship. |
| Edge disruption score | $d_{x_{ij}}^{(E)}$ | $d_{x_{ij}}^{(E)} = \lvert E^*_{x_{ij}} - B^*_{ij} \rvert$ | Edge disruption scores measure the extent to which a particular TF-gene regulatory relationship is disrupted by genetic variants in a given individual. A high value of $d_{x_{ij}}^{(E)}$ indicates that the regulatory relationship between TF $i$ and gene $j$ is likely disrupted by genetic variants. |
| TF disruption score | $d_{x_i}^{(TF)}$ | $d_{x_i}^{(TF)} = \sum_j \left\lvert E^*_{x_{ij}} - B^*_{ij} \right\rvert$ | TF disruption scores measure the extent to which a particular TF's binding sites in promoters across the genome are disrupted by genetic variants in a given individual. A high value of $d_{x_i}^{(TF)}$ indicates that the binding sites of TF $i$ are likely disrupted by genetic variants. |
| Gene disruption score | $d_{x_j}^{(G)}$ | $d_{x_j}^{(G)} = \sum_i \left\lvert E^*_{x_{ij}} - B^*_{ij} \right\rvert$ | Gene disruption scores measure extent to which a particular gene's promoter region is disrupted by genetic variants in a given individual. A high value of $d_{x_j}^{(G)}$ indicates that the promoter region of gene $j$ is likely disrupted by genetic variants. |
| Edge regulatory difference score | $R_{ij}^{(E)}$ | $R_{ij}^{(E)} = \left\lvert d_{g_{ij}}^{(E)} - d_{k_{ij}}^{(E)} \right\rvert$ | Edge regulatory difference scores compare the edge disruption scores between two individuals, and thus measure the extent to which a TF-gene relationship is differentially disrupted by genetic variants between two individuals. |

| | | | |
|---|---|---|---|
| Gene regulatory difference score | $R_j^{(G)}$ | $R_j^{(G)} = \sum_i R_{ij}^{(E)}$ | Gene regulatory difference scores sum the edge regulatory difference scores per gene, and thus measure the extent to which the promoter region of a gene is differentially disrupted by genetic variants when comparing two individuals. |
| Differential modularity score | $DM$ | ALPACA[23] | ALPACA, when applied to compare an EGRET GRN $E^*$ with a baseline $B^*$, calculates a differential modularity score for each gene. The DM score indicates the contribution of that gene to the differential modularity between the baseline and EGRET GRNs. |

Table S3: Improvement in AUC-ROC for the prediction of the ChIP-seq regulatory network in GM12878 when using EGRET edge weights, over using baseline network edge-weights, for different cutoffs of $d_{x_{ij}}^{(E)}$. Total number of negatives (N), total number of positives (P), improvement in the AUC-ROC as well as the Delong p-value for the improvement are reported.

| $d_{x_{ij}}^{(E)}$ **cutoff** | **N** | **P** | **AUC improvement** | **Delong p-value** |
|---|---|---|---|---|
| 0.1 | 226 | 133 | -0.05 | 0.96 |
| 0.15 | 132 | 81 | -0.07 | 0.95 |
| 0.2 | 90 | 76 | -0.01 | 0.55 |
| 0.25 | 72 | 75 | 0.08 | 0.07 |
| 0.3 | 70 | 72 | 0.09 | 0.05 |
| **0.35** | **57** | **65** | **0.14** | **0.01** |
| 0.4 | 57 | 64 | 0.13 | 0.02 |
| 0.45 | 57 | 64 | 0.13 | 0.02 |
| 0.5 | 57 | 64 | 0.13 | 0.02 |
| 0.55 | 57 | 62 | 0.11 | 0.04 |
| 0.6 | 57 | 62 | 0.11 | 0.04 |
| 0.65 | 57 | 61 | 0.10 | 0.05 |
| 0.7 | 57 | 61 | 0.10 | 0.05 |
| 0.75 | 57 | 58 | 0.07 | 0.12 |
| 0.8 | 57 | 58 | 0.07 | 0.12 |
| 0.85 | 57 | 58 | 0.07 | 0.12 |
| 0.9 | 57 | 57 | 0.06 | 0.16 |
| 1 | 56 | 56 | 0.06 | 0.16 |

Table S4: Improvement in AUC-ROC for the prediction of the ChIP-seq regulatory network in K562 when using EGRET edge weights, over using baseline network edge-weights, for different cutoffs of $d_{x_{ij}}^{(E)}$. Total number of negatives (N), total number of positives (P), improvement in the AUC-ROC as well as the Delong p-value for the improvement are reported.

| $d_{x_{ij}}^{(E)}$ cutoff | N | P | AUC improvement | Delong p-value |
|---|---|---|---|---|
| 0.1 | 750 | 547 | -0.01 | 0.88 |
| 0.15 | 408 | 283 | -0.03 | 0.90 |
| 0.2 | 235 | 161 | -0.05 | 0.93 |
| 0.25 | 149 | 127 | -0.01 | 0.55 |
| 0.3 | 105 | 97 | 0.03 | 0.29 |
| **0.35** | **75** | **78** | **0.11** | **0.03** |
| 0.4 | 68 | 72 | 0.14 | 0.01 |
| 0.45 | 67 | 70 | 0.13 | 0.02 |
| 0.5 | 67 | 69 | 0.13 | 0.02 |
| 0.55 | 67 | 68 | 0.12 | 0.03 |
| 0.6 | 67 | 68 | 0.12 | 0.03 |
| 0.65 | 64 | 63 | 0.12 | 0.03 |
| 0.7 | 61 | 57 | 0.11 | 0.05 |
| 0.75 | 61 | 57 | 0.11 | 0.05 |
| 0.8 | 61 | 57 | 0.11 | 0.05 |
| 0.85 | 61 | 57 | 0.11 | 0.05 |
| 0.9 | 61 | 57 | 0.11 | 0.05 |
| 1 | 61 | 56 | 0.11 | 0.05 |

Table S5: GWAS catalog study references for CAD genes.

| PMID | First author | Date | Journal | Study | Ref |
|---|---|---|---|---|---|
| 21239051 | Reilly MP | 2011-01-14 | Lancet | Identification of ADAMTS7 as a novel locus for coronary atherosclerosis and association of ABO with myocardial infarction in the presence of coronary atherosclerosis: two genome-wide association studies. | [26] |
| 24262325 | Dichgans M | 2013-11-21 | Stroke | Shared genetic susceptibility to ischemic stroke and coronary artery disease: a genome-wide analysis of common variants. | [27] |
| 26343387 | Nikpay M | 2015-09-07 | Nat Genet | A comprehensive 1,000 Genomes-based genome-wide association meta-analysis of coronary artery disease. | [28] |
| 26708285 | Wakil SM | 2016-02-01 | Atherosclerosis | A genome-wide association study reveals susceptibility loci for myocardial infarction/coronary artery disease in Saudi Arabs. | [29] |
| 28714974 | Klarin D | 2017-07-17 | Nat Genet | Genetic analysis in UK Biobank links insulin resistance and transendothelial migration pathways to coronary artery disease. | [30] |
| 29212778 | van der Harst P | 2017-12-06 | Circ Res | Identification of 64 Novel Genetic Loci Provides an Expanded View on the Genetic Architecture of Coronary Artery Disease. | [31] |
| 29263402 | Han Y | 2017-12-20 | Sci Rep | Genome-wide association study identifies a missense variant at APOA5 for coronary artery disease in Multi-Ethnic Cohorts from Southeast Asia. | [32] |
| 29472232 | Li Y | 2018-02-22 | Arterioscler Thromb Vasc Biol | Genome-Wide Association and Functional Studies Identify SCML4 and THSD7A as Novel Susceptibility Genes for Coronary Artery Disease. | [33] |
| 30104761 | Zhou W | 2018-08-13 | Nat Genet | Efficiently controlling for case-control imbalance and sample relatedness in large-scale genetic association studies. | [34] |
| 30402224 | Yamada Y | 2018-09-17 | Biomed Rep | Identification of 26 novel loci that confer susceptibility to early-onset coronary artery disease in a Japanese population. | [35] |

Table S6: GWAS catalog study references for CD genes.

| PMID | Journal | Study | Ref |
|---|---|---|---|
| 17435756 | Nat Genet | Genome-wide association study identifies new susceptibility loci for Crohn disease and implicates autophagy in disease pathogenesis. | [36] |
| 17447842 | PLoS Genet | Novel Crohn disease locus identified by genome-wide association maps to a gene desert on 5p13.1 and modulates expression of PTGER4. | [37] |
| 17554261 | Nat Genet | Sequence variants in the autophagy gene IRGM and multiple other replicating loci contribute to Crohn's disease susceptibility. | [38] |
| 17554300 | Nature | Genome-wide association study of 14,000 cases of seven common diseases and 3,000 shared controls. | [39] |
| 17684544 | PLoS One | Systematic association mapping identifies NELL1 as a novel IBD disease gene. | [40] |
| 17804789 | Proc Natl Acad Sci U S A | Genome-wide association study for Crohn's disease in the Quebec Founder Population identifies multiple validated disease loci. | [41] |
| 18587394 | Nat Genet | Genome-wide association defines more than 30 distinct susceptibility loci for Crohn's disease. | [42] |
| 20570966 | Hum Mol Genet | Fucosyltransferase 2 (FUT2) non-secretor status is associated with Crohn's disease. | [43] |
| 21102463 | Nat Genet | Genome-wide meta-analysis increases to 71 the number of confirmed Crohn's disease susceptibility loci. | [44] |
| 22293688 | Eur J Hum Genet | 1000 Genomes-based imputation identifies novel and refined associations for the Wellcome Trust Case Control Consortium phase 1 Data. | [45] |
| 22412388 | PLoS Genet | A genome-wide scan of Ashkenazi Jewish Crohn's disease suggests novel susceptibility loci. | [46] |
| 22936669 | Gut | A genome-wide association study on a southern European population identifies a new Crohn's disease susceptibility locus at RBX1-EP300. | [47] |
| 23128233 | Nature | Host-microbe interactions have shaped the genetic architecture of inflammatory bowel disease. | [48] |
| 23266558 | Gastroenterology | A genome-wide association study identifies 2 susceptibility Loci for Crohn's disease in a Japanese population. | [49] |
| 23850713 | Gut | Genome-wide association study of Crohn's disease in Koreans revealed three new susceptibility loci and common attributes of genetic susceptibility across ethnic populations. | [50] |
| 25489960 | Inflamm Bowel Dis | Immunochip analysis identification of 6 additional susceptibility loci for Crohn's disease in Koreans. | [51] |
| 26192919 | Nat Genet | Association analyses identify 38 susceptibility loci for inflammatory bowel disease and highlight shared genetic risk across populations. | [52] |
| 26278503 | Gastroenterology | Characterization of genetic loci that affect susceptibility to inflammatory bowel diseases in African Americans. | [53] |
| 26891255 | Inflamm Bowel Dis | HLA-C*01 is a Risk Factor for Crohn's Disease. | [54] |
| 28008999 | Sci Rep | Genetic architecture differences between pediatric and adult-onset inflammatory bowel diseases in the Polish population. | [55] |
| 28067908 | Nat Genet | Genome-wide association study implicates immune activation of multiple integrin genes in inflammatory bowel disease. | [56] |
| 30500874 | J Crohns Colitis | A genome-wide association study identifying RAP1A as a novel susceptibility gene for Crohn's disease in Japanese individuals. | [57] |

Table S7: GO terms enriched in the genes with highest DM scores in individual 18 (genotype NA18523) in CMs. Enrichment performed using GORILLA [24]. N - total number of genes, B - total number of genes associated with a specific GO term, n - number of genes in the top of the user's input list, b - number of genes in the intersection

| GO Term | Description | P-value FDR | q-value | Enrichment | N | B | n | b | Genes |
|---|---|---|---|---|---|---|---|---|---|
| *Table available at* https://github.com/daweighill/EGRET/tree/master/supplementary_tables | | | | | | | | | |

Table S8: GO terms enriched in the genes with highest DM scores in individual 18 (genotype NA18523) in LCLs. Enrichment performed using GORILLA [24]. N - total number of genes, B - total number of genes associated with a specific GO term, n - number of genes in the top of the user's input list, b - number of genes in the intersection

| GO Term | Description | P-value FDR | q-value | Enrichment | N | B | n | b | Genes |
|---|---|---|---|---|---|---|---|---|---|
| *Table available at* https://github.com/daweighill/EGRET/tree/master/supplementary_tables | | | | | | | | | |

Table S9: GO terms enriched in the genes with highest DM scores in individual 18 (genotype NA18523) in iPSCs. Enrichment performed using GORILLA [24]. N - total number of genes, B - total number of genes associated with a specific GO term, n - number of genes in the top of the user's input list, b - number of genes in the intersection

| GO Term | Description | P-value FDR | q-value | Enrichment | N | B | n | b | Genes |
|---|---|---|---|---|---|---|---|---|---|
| *Table available at* https://github.com/daweighill/EGRET/tree/master/supplementary_tables | | | | | | | | | |

#### References


[1] Charles E Grant, Timothy L Bailey, and William Stafford Noble. Fimo: scanning for occurrences of a given motif. *Bioinformatics*, 27(7):1017–1018, 2011.

[2] Michael Lawrence, Wolfgang Huber, Hervé Pagès, Patrick Aboyoun, Marc Carlson, Robert Gentleman, Martin Morgan, and Vincent Carey. Software for computing and annotating genomic ranges. *PLoS Computational Biology*, 9, 2013.

[3] John Lonsdale, Jeffrey Thomas, Mike Salvatore, Rebecca Phillips, Edmund Lo, Saboor Shad, Richard Hasz, Gary Walters, Fernando Garcia, Nancy Young, et al. The genotype-tissue expression (GTEx) project. *Nature genetics*, 45(6):580, 2013.

[4] Bo Zhou, Steve S Ho, Stephanie U Greer, Xiaowei Zhu, John M Bell, Joseph G Arthur, Noah Spies, Xianglong Zhang, Seunggyu Byeon, Reenal Pattni, et al. Comprehensive, integrated, and phased whole-genome analysis of the primary encode cell line k562. *Genome research*, 29(3):472–484, 2019.

[5] Vincentius Martin, Jingkang Zhao, Ariel Afek, Zachery Mielko, and Raluca Gordân. Qbic-pred: quantitative predictions of transcription factor binding changes due to sequence variants. *Nucleic acids research*, 47(W1):W127–W135, 2019.

[6] Abhijeet Rajendra Sonawane, John Platig, Maud Fagny, Cho-Yi Chen, Joseph Nathaniel Paulson, Camila Miranda Lopes-Ramos, Dawn Lisa DeMeo, John Quackenbush, Kimberly Glass, and Marieke Lydia Kuijjer. Understanding tissue-specific gene regulation. *Cell reports*, 21(4):1077–1088, 2017.

[7] Kimberly Glass, Curtis Huttenhower, John Quackenbush, and Guo-Cheng Yuan. Passing messages between biological networks to refine predicted interactions. *PloS one*, 8(5):e64832, 2013.

[8] Kimberly Glass, John Quackenbush, Edwin K Silverman, Bartolome Celli, Stephen I Rennard, Guo-Cheng Yuan, and Dawn L DeMeo. Sexually-dimorphic targeting of functionally-related genes in copd. *BMC systems biology*, 8(1):1–17, 2014.

[9] Weiliang Qiu, Feng Guo, Kimberly Glass, Guo Cheng Yuan, John Quackenbush, Xiaobo Zhou, and Kelan G Tantisira. Differential connectivity of gene regulatory networks distinguishes corticosteroid response in asthma. *Journal of Allergy and Clinical Immunology*, 141(4):1250–1258, 2018.

[10] Kimberly Glass, John Quackenbush, Dimitrios Spentzos, Benjamin Haibe-Kains, and Guo-Cheng Yuan. A network model for angiogenesis in ovarian cancer. *BMC bioinformatics*, 16(1):1–17, 2015.

[11] Camila M Lopes-Ramos, Marieke L Kuijjer, Shuji Ogino, Charles S Fuchs, Dawn L DeMeo, Kimberly Glass, and John Quackenbush. Gene regulatory network analysis identifies sex-linked differences in colon cancer drug metabolism. *Cancer research*, 78(19):5538–5547, 2018.

[12] Camila M Lopes-Ramos, Cho-Yi Chen, Marieke L Kuijjer, Joseph N Paulson, Abhijeet R Sonawane, Maud Fagny, John Platig, Kimberly Glass, John Quackenbush, and Dawn L DeMeo. Sex differences in gene expression and regulatory networks across 29 human tissues. *Cell reports*, 31(12):107795, 2020.

[13] Jeanne Chèneby, Marius Gheorghe, Marie Artufel, Anthony Mathelier, and Benoit Ballester. Remap 2018: an updated atlas of regulatory regions from an integrative analysis of dna-binding chip-seq experiments. *Nucleic acids research*, 46(D1):D267–D275, 2018.

[14] Takaya Saito and Marc Rehmsmeier. Precrec: fast and accurate precision-recall and roc curve calculations in r. *Bioinformatics*, 33 (1):145–147, 2017.

[15] Xavier Robin, Natacha Turck, Alexandre Hainard, Natalia Tiberti, Frédérique Lisacek, Jean-Charles Sanchez, and Markus Müller. proc: an open-source package for r and s+ to analyze and compare roc curves. *BMC Bioinformatics*, 12:77, 2011.

[16] Cynthia A Kalita, Christopher D Brown, Andrew Freiman, Jenna Isherwood, Xiaoquan Wen, Roger Pique-Regi, and Francesca Luca. High-throughput characterization of genetic effects on dna–protein binding and gene transcription. *Genome research*, 28(11):1701–1708, 2018.

[17] Nicholas E Banovich, Yang I Li, Anil Raj, Michelle C Ward, Peyton Greenside, Diego Calderon, Po Yuan Tung, Jonathan E Burnett, Marsha Myrthil, Samantha M Thomas, et al. Impact of regulatory variation across human ipscs and differentiated cells. *Genome research*, 28(1):122–131, 2018.

[18] Yang I Li, Bryce Van De Geijn, Anil Raj, David A Knowles, Allegra A Petti, David Golan, Yoav Gilad, and Jonathan K Pritchard. Rna splicing is a primary link between genetic variation and disease. *Science*, 352(6285):600–604, 2016.

[19] Jacob F Degner, Athma A Pai, Roger Pique-Regi, Jean-Baptiste Veyrieras, Daniel J Gaffney, Joseph K Pickrell, Sherryl De Leon, Katelyn Michelini, Noah Lewellen, Gregory E Crawford, et al. Dnase i sensitivity qtls are a major determinant of human expression variation. *Nature*, 482(7385):390–394, 2012.

[20] Benjamin M Bolstad, Rafael A Irizarry, Magnus Åstrand, and Terence P. Speed. A comparison of normalization methods for high density oligonucleotide array data based on variance and bias. *Bioinformatics*, 19(2):185–193, 2003.

[21] Ole Tange. Gnu parallel-the command-line power tool.; login: The usenix magazine, 36 (1): 42–47, 2011.

[22] Annalisa Buniello, Jacqueline A L MacArthur, Maria Cerezo, Laura W Harris, James Hayhurst, Cinzia Malangone, Aoife McMahon, Joannella Morales, Edward Mountjoy, Elliot Sollis, et al. The nhgri-ebi gwas catalog of published genome-wide association studies, targeted arrays and summary statistics 2019. *Nucleic acids research*, 47(D1):D1005–D1012, 2019.
